# Predicting Early Diabetic Kidney Disease Using a 3D Mesangial Model and Bayesian Mechano-Chemical Stratification

**DOI:** 10.1101/2023.12.10.571017

**Authors:** Biswajoy Ghosh, Farhad Niknam, Nikhil Kumar, Kristin Andreassen Fenton, Krishna Agarwal

**Affiliations:** Department of Physics and Technology, UiT-The Arctic University of Norway, Tromsø, Norway; Advanced Technology Development Centre, Indian Institute of technology Kharagpur, India; Department of Medical Biology, UiT-The Arctic University of Norway, Tromsø, Norway

## Abstract

Diabetic kidney disease (DKD) is often diagnosed only after irreversible damage, limiting early treatment. The kidney mesangium, a structure highly sensitive to mechanical forces, plays a key role in early fibrosis development, yet its response to early disease cues like stiffness and biochemical changes is poorly understood. This is due to limitations in current in vitro models, poor in vivo accessibility, and static biopsy samples. To address this, we developed a 3D in vitro model of the mesangial microenvironment using stiffness-tunable gelatin methacrylate (GelMA) hydrogels that mimic healthy and fibrotic kidney tissue. Exposing mesangial cells to glucose and TGF-*β*1 led to altered cell shape, increased dry mass, and elevated expression of fibrotic markers (*α*-SMA and collagen IV), especially under stiffer conditions, indicating a synergistic effect of biochemical and mechanical stress. These responses were integrated using Gaussian process regression to create a 3D “severity cube” that maps DKD progression across mechanical and chemical inputs. This system quantifies early mesangial transitions and reveals key fibrosis-related mechanisms. By combining organotypic modeling with interpretable inference, our platform offers a predictive tool for early disease stratification and a basis for studying subclinical fibrosis progression in DKD.

## INTRODUCTION

Diabetic kidney disease (DKD) is one of the most common causes of end-stage renal failure, affecting nearly one-third of people with diabetes worldwide^1^. Despite its prevalence and clinical burden, DKD is typically diagnosed only after substantial nephron damage has already occurred. This diagnostic delay severely limits treatment options, as current interventions can only slow progression rather than reverse pathology once fibrotic remodeling has set in. Standard clinical markers—such as albuminuria and estimated glomerular filtration rate (eGFR)—are often insufficient to capture the earliest stages of disease, underscoring an urgent need for predictive tools that can detect subclinical transitions before irreversible injury. At the core of this unmet need lies a biological blind spot: the early changes in cellular behavior and microenvironmental remodeling that precede the obvious structural damage. Among the key players in this process are the mesangial cells, located within the kidney’s glomerular filtration unit. Mesangial cells are uniquely poised to sense and integrate biochemical and mechanical stimuli. They are also among the first to respond to diabetic stressors, such as hyperglycemia and inflammatory cytokines. In healthy tissue, mesangial cells regulate capillary tone, clear debris, and maintain extracellular matrix (ECM) homeostasis^2–6^. In DKD, however, they contribute to mesangial expansion through increased proliferation and excessive secretion of ECM components like collagen IV, fibronectin, and laminins^7,8^. This fibrogenic activity not only thickens the glomerular basement membrane but also increases matrix stiffness—an underappreciated driver of disease progression.

Importantly, the mesangial microenvironment is both chemically and mechanically active. Elevated glucose and cytokines such as TGF-*β* are well-established biochemical triggers of fibrosis^4,9,10^, but increasing matrix stiffness itself can reinforce fibrogenic signaling. This suggests that DKD may progress through a nonlinear interplay between chemical and mechanical cues^11^. Yet, our ability to unravel these early-stage transitions has been limited by a lack of appropriate model systems. Conventional 2D cultures oversimplify cellular context and fail to replicate mechanical realism; animal models obscure causality; and human biopsy samples represent only late-stage, static disease states. Consequently, a mechanistic understanding of how mesangial cells integrate mechano-chemical inputs—and how these inputs might predict disease trajectory—remains elusive^12^.

To fill this gap, we propose a predictive, non-invasive diagnostics for DKD requires experimental systems that recapitulate the biophysical microenvironment of early disease. Such systems must support high-resolution readouts of mesangial response while allowing for systematic variation of matrix stiffness and chemical stimuli. Moreover, they must be capable of producing interpretable, quantitative frameworks that relate cellular behavior to disease severity. Recent advances in engineered biomaterials and data-driven modeling provide an opportunity to achieve this integration.

In this study, we present a 3D in vitro model of the mesangial niche that enables interrogation of stiffness- and cytokine-driven transitions relevant to DKD. Using tunable gelatin methacrylate (GelMA) hydrogels to emulate the stiffness range from healthy to fibrotic mesangial matrix, we expose mesangial cells to defined mechano-chemical conditions and assess their morphological, molecular, and biosynthetic responses. To translate these responses into a predictive framework, we introduce a probabilistic model using the Gaussian process regression method trained on fibrosis-relevant markers^13^, *α*-smooth muscle actin (*α*-SMA) and collagen IV, to construct a continuous severity cube. This model captures disease progression across a spectrum of mechanical and biochemical stimuli and supports interpolation of intermediate states beyond those directly tested. By bridging high-fidelity modeling of the mesangial microenvironment with a probabilistic, interpretable framework for disease stratification, this work provides a foundational step toward predictive diagnostics for DKD. It also offers a generalizable strategy for modeling early pathophysiological transitions in other mechano-responsive diseases.

## RESULTS

### Evaluating 3D In-Vitro Mesangium Model to Match Healthy and Fibrotic Kidney Mesangium

We made and evaluated 3D hydrogels with different GelMA concentrations. We found that 5% GelMA closely replicated the mechanical properties of healthy mesangium (0.4-3 kPa)^14^. 10% GelMA on the other hand demonstarted about 8-10 × higher stiffness tyoical of fibrotic mesangium (8-30 kPa)^15^ in diabetic nephropathy^16^. Additionally, for these concentrations, we conducted multimodal mechanical characterization to evaluate the physical similarity of the 3D gels with mesangial tissue and promote reproducibility.

A schematic (Figure 1a) illustrates the shift in mesangial stiffness during nephropathy progression and our 3D in vitro model approach incorporating mesangial cells in controlled stiffness and chemical conditions. Compression tests (Figure 1b–c) confirmed that 10% GelMA hydrogels exhibit higher compressive stress than 5% counterparts (8-10 times higher), validating increased bulk stiffness representative of fibrotic mesangium. Shear-dependent viscosity and stress measurements (Figure 1d–f) revealed that both 5% and 10% hydrogels exhibit shear-thinning behavior typical of viscoelastic materials like tissues, with 10% GelMA showing higher viscosity and increased resistance to shear, indicating elevated viscoelastic integrity. Figure 1f indicates that the 10% GelMA hydrogels exhibit increased shear stress across shear rates, suggesting a denser polymeric network and enhanced resistance to deformation under mechanical loading due to higher polymer concentration and crosslink density, closely resembling the stiff, fibrotic mesangial environment in diabetic kidney disease^15^; conversely, the lower shear stress of 5% GelMA aligns with the softer mechanical characteristics typical of healthy mesangial tissue. Rheological assessment across physiological temperatures (Figure 1g–h) indicated that 10% GelMA retained a higher storage modulus than 5%, with minimal fluctuation, confirming mechanical stability. Dynamic mechanical analysis under varying frequencies (Figure 1i–j) further demonstrated a stiffer response in 10% GelMA across all tested frequencies. Thermal analysis by differential scanning calorimetry (Figure 1k) revealed a more pronounced endothermic transition in 10% GelMA, supporting increased crosslink density. A detailed chemical and mechanical analysis was performed to ensure reproducibility and consistency in the hydrogels (Refer to the supplementary Figures S1-S4).

**Figure 1.**
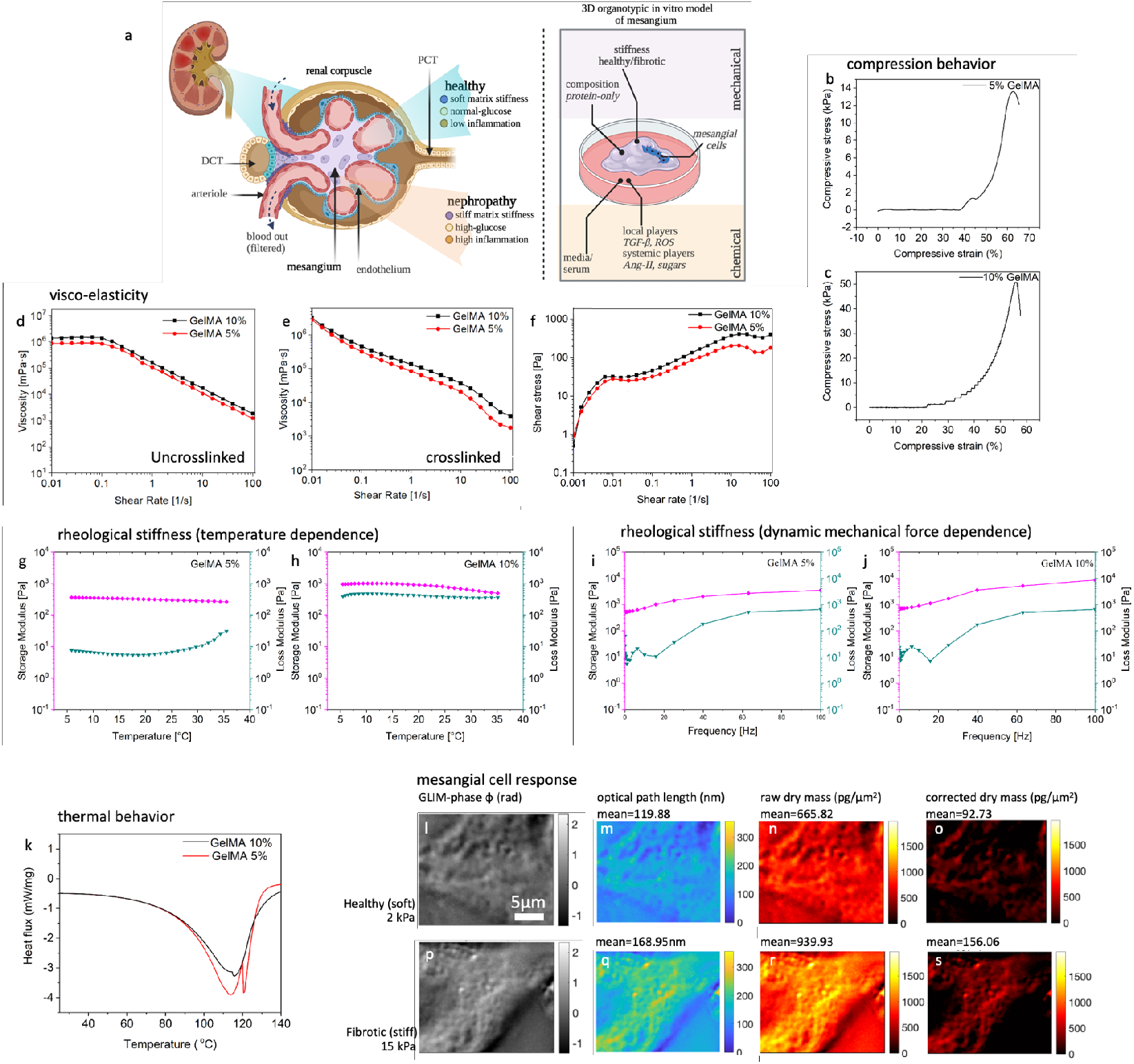
Development of a 3D in vitro organotypic model of the kidney mesangium with tunable mechanical microenvironments mimicking healthy and fibrotic states. (a) Schematic of kidney glomerular filtration unit, highlighting the mesangium in healthy (soft, low inflammation) and fibrotic (stiff, inflamed) states in diabetic nephropathy. (b–c) Compression analysis of 5% GelMA shows physiological stiffness and 10% GelMA hydrogels shows about 8-10 ×higher stiffness typical of fibrotic tissues. (d) Viscosity–shear rate profiles of uncrosslinked 5% GelMA hydrogels show shear-thinning behavior, with higher viscosity in 10% GelMA, indicating denser polymer entanglement prior to crosslinking. (e) In crosslinked hydrogels, 10% GelMA retains consistently higher viscosity across all shear rates, reflecting increased internal friction and viscoelastic damping typical of fibrotic tissue. (f) Shear stress measurements confirm that 10% GelMA exhibits higher resistance to shear deformation, indicating increased crosslink density and mechanical stiffness.(g–h) Temperature-dependent rheological analysis shows that 10% GelMA hydrogels maintain a consistently higher storage modulus (G) across physiological temperatures, indicating greater mechanical stability and stiffness compared to 5% GelMA. (i–j) Frequency sweep rheology further underscores the mechanical contrast, with both storage and loss moduli increasing with GelMA concentration and frequency, confirming the enhanced viscoelastic stiffness of fibrotic-like matrices.(k) Differential scanning calorimetry (DSC) reveals a more prominent endothermic transition in 10% GelMA, indicating greater crosslink density and thermal stability compared to 5% GelMA. (l–s) Gradient Light Interference Microscopy (GLIM) images showing the morphology and dry mass distribution of mesangial cells cultured on soft (5%, healthy-like) and stiff (10%, fibrotic-like) hydrogels. Cells on fibrotic matrices exhibit denser structures and higher intracellular dry mass, suggesting matrix stiffness–dependent cellular adaptations in growth and biosynthesis.

High-resolution gradient light interference microsocpy (GLIM) imaging (Figure 1l–s) were used to visualize cell-hydrogel interactions at high resolution and label-free^17,18^. GLIM enabled quantitative measurement of cellular dry mass and optical thickness which measures non-water components like protein levels and cytoskeletal integrity^19,20^. Cells grown on stiffer (10%) GelMA matrices displayed higher dry mass content and denser intracellular profiles compared to those on soft (5%) matrices (Figure 1l–s). Background effects due to substrate were subtracted to get the true value of dry mass in the cells. The supplementary Figure S5 shows both larger fields in the 5% and 10% GelMA as well as nucleo-cytoplasmic density changes. These findings indicate that matrix stiffness influences cell biosynthetic activity of mesangial expansion, reinforcing the physiological relevance of the hydrogel system in mimicking healthy versus fibrotic mesangial environments^21^.

These results establish the ability of our engineered hygrogel-based in vitro system to mimic the mechanical landscape of both healthy and diseased mesangial environments, forming the foundation for further biological studies in diabetic kidney disease models.

### Mesangial Cell Morphodynamics Are Governed by Matrix Stiffness and TGF-*β* 1-Induced Mechano-Chemical Cues in High Glucose Environment

To investigate how mechano-chemical cues influence mesangial cell morphology, we cultured cells in four conditions combining matrix stiffness (healthy vs. fibrotic) with biochemical stimulation (± TGF-*β* 1). Representative images (Figure 2a–d) show that matrix stiffness and TGF-*β* 1 alter cellular morphology. Cells on healthy matrices displayed an elongated, spread-out phenotype with fine projections (Figure 2f), while fibrotic matrices induced compact, contracted shapes (Figure 2h–i). TGF-*β* 1 further exaggerated this shift, promoting rounded, clustered morphologies, especially in stiff matrices (Figure 2b,d). Quantitative shape analysis confirmed a significant increase in compactness with increasing matrix stiffness and TGF-*β* 1 treatment (Figure 2e), indicating a shift toward stronger cell–cell interactions over cell–matrix interactions. Compactness here means the quantification of how densely mesangial cells cluster together, measured as the percentage of cell-covered area relative to gaps between cells. Higher compactness indicates stronger cell–cell interactions and tighter clustering, characteristic of fibrotic environments. Filopodia length analysis (Figure 2j) revealed longer, more exploratory extensions in healthy matrix conditions. This was reduced significantly in fibrotic and TGF-*β* 1-treated groups, suggesting compromised actin-driven protrusive behavior. Time-lapse imaging of filopodia (Figure 2k) further demonstrated that mesangial cells on healthy matrices maintain persistent and dynamic filopodia, while cells on fibrotic matrices exhibited more static, bundled protrusions. Directional plasticity, quantified via angular projection analysis (Figure 2l), was significantly lower in the fibrotic matrix, supporting a transition toward more directed, less exploratory migration behavior. These results demonstrate that mesangial cell morphology and plasticity are finely regulated by the mechanical and chemical composition of the microenvironment^22^, with fibrotic-like conditions reducing cellular exploratory behavior and enhancing compaction, a hallmark of fibrosis progression^13^.

**Figure 2.**
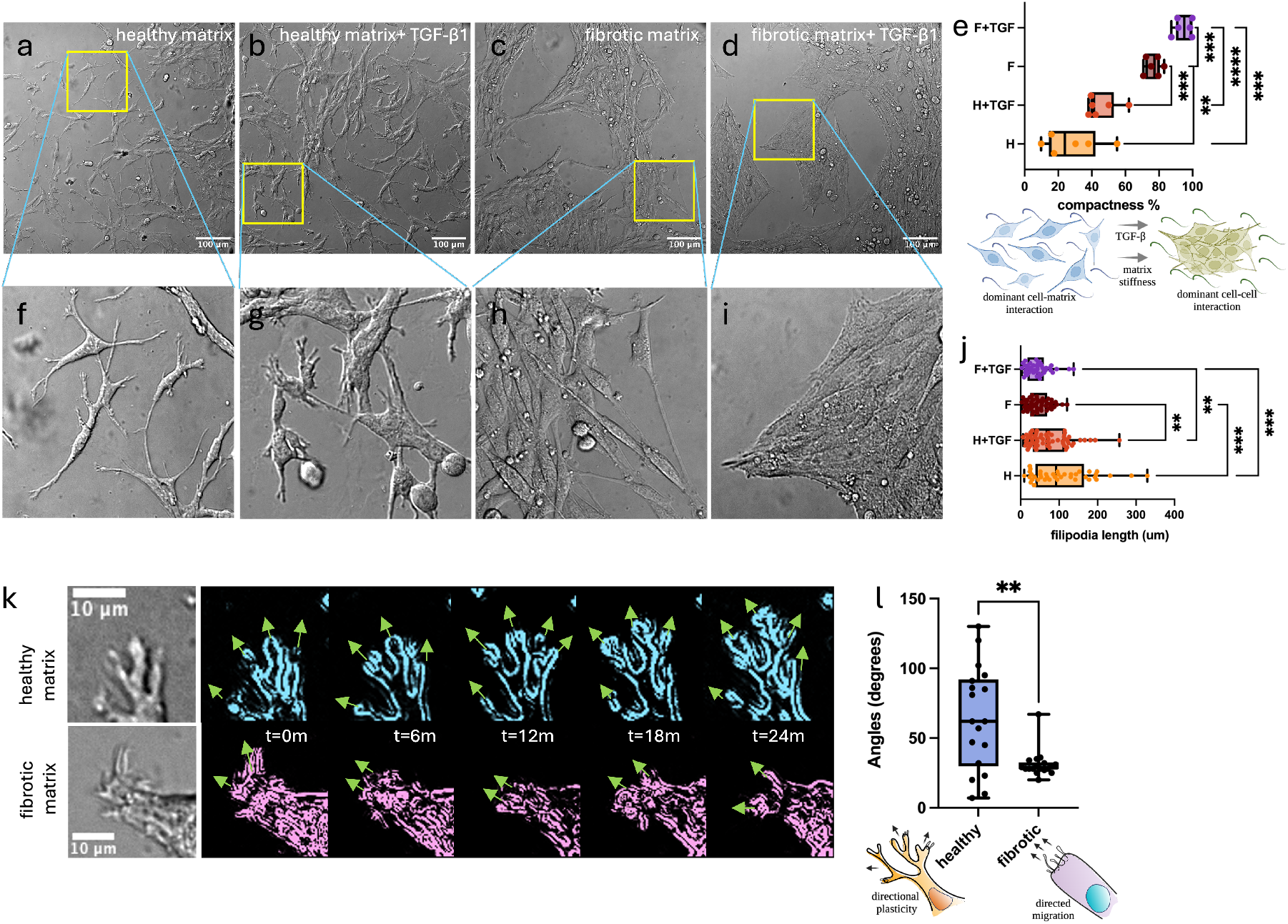
Mesangial cell morphology is modulated by matrix stiffness and TGF-*β* 1 signaling, revealing mechano-chemical control over cell plasticity. (a–d) Brightfield microscopy images of mesangial cells cultured on: (a) healthy matrix, (b) healthy matrix + TGF-*β* 1, (c) fibrotic matrix, (d) fibrotic matrix + TGF-*β* 1. Scale bars: 100 *µ*m. (f–i) High-magnification views of yellow-boxed regions in panels (a–d), revealing changes in cell shape and spreading. (e) Quantification of cell compactness shows increased compactness in fibrotic conditions, especially with TGF-*β* 1. (j) Quantification of filopodia length reveals longer and branched filopodia in healthy matrix conditions and reduced protrusions in fibrotic matrix ± TGF-*β* 1. (k) Time-lapse imaging and segmentation of filopodia morphology show persistent, dynamic protrusions in healthy matrix and stunted, less plastic behavior in fibrotic matrix over 24 minutes (arrows indicate directionality). (l) Angular spread analysis of filopodial projections demonstrates decreased directional plasticity in fibrotic matrix conditions. Schematic illustrates the shift from exploratory morphology in healthy states to directed morphology in fibrosis. Statistical test done with 2-way ANOVA, multiple comparisons between each category is done, only significant groups highlighted in the plot. ns (not significant): 0.123, *: p 0.032, **: 0021, ***: 0.0002, ****: <0.0001

Moreover, our detailed experiments showed that mesangial cells cultured on glass surfaces show faster proliferation and a lamellar morphology, especially under high glucose conditions, whereas cells on the GelMA matrix demonstrate regulated proliferation and maintain a branched, process-rich morphology typical of mesangial cells (shown in supplementary Figure S6. This is because the glass substrate, unlike the 3D in-vitro mesangium model, is hard and does not emulate the chemical and mechanical environment that of the soft human tissues. This difference stresses the importance of the substrate’s mechanical and chemical properties in emulating in vivo conditions.

### Matrix Stiffness Amplifies Glucose- and TGF-*β* 1-Induced Fibrosis in 3D Mesangium Model of Diabetic Kidney Disease

The schematic (Figure 3d) illustrates the role of the chemical (glucose, TGF) and mechanical stressors (matrix stiffenss) in DKD progression. The influence of the stressors in isolation as well as together on pathogenesis can be quantified by expression levels of *α*-SMA and Collagen-IV, as key biomarkers of DKD severity^23^. High glucose and inflammatory cues activate *α*-SMA and Collagen IV expression, with matrix stiffness acting as an amplifier that accelerates mesangial cell transition toward a fibrotic phenotype. The increased expression of the pathological markers with the DKD stressors, suggest relevance of our in vitro 3D mesangium model to capture the DKD progression. These markers were assessed under a range of mechano-chemical conditions that simulate different stages of diabetic nephropathy. *α* − SMA is strongly correlated to cellular contractility typical in DKD^24^. Cells were cultured on control (glass), healthy-like, or fibrotic-like GelMA matrices under low or high glucose concentrations, with or without TGF-*β* 1 (3a-c). It was observed that *α*-SMA expression was low in healthy matrix under low glucose conditions but increased significantly (1.5 to 2 times) with either TGF-*β* 1 treatment or high glucose (Figure 3b, e). The highest *α*-SMA expression was observed in cells on stiff matrices exposed to both high glucose and TGF-*β* 1 (Figure 3c), indicating a synergistic effect between matrix stiffness and inflammatory/glucose signals. Quantitative analysis (Figure 3e, supplementary Table 1) confirmed a statistically significant increase in *α*-SMA intensity in stiff hydrogels compared to control or healthy matrix. Collagen IV is the key matrix protein that the mesangial cells deposit and contribute to the mesangium and the basement membrane^25^. Collagen IV deposition (Figure 3f–h) was minimal under basal conditions but increased dramatically in fibrotic matrices and further amplified with high glucose and TGF-*β* 1. The Box plots (Figure 3i–j) indicated significant increases in Collagen IV under both low and high glucose conditions, but with steeper upregulation on fibrotic matrices, illustrating the strong role of mechanical stiffness in ECM remodeling. Detailed spatial analysis and further quantification of fibrotic markers under various mechano-chemical conditions are provided in supplementary Figures S7–S9. This leads to thickening of the basement membrane, reduced glomerular capillary flexibility, and disrupted intercellular communication.

**Figure 3.**
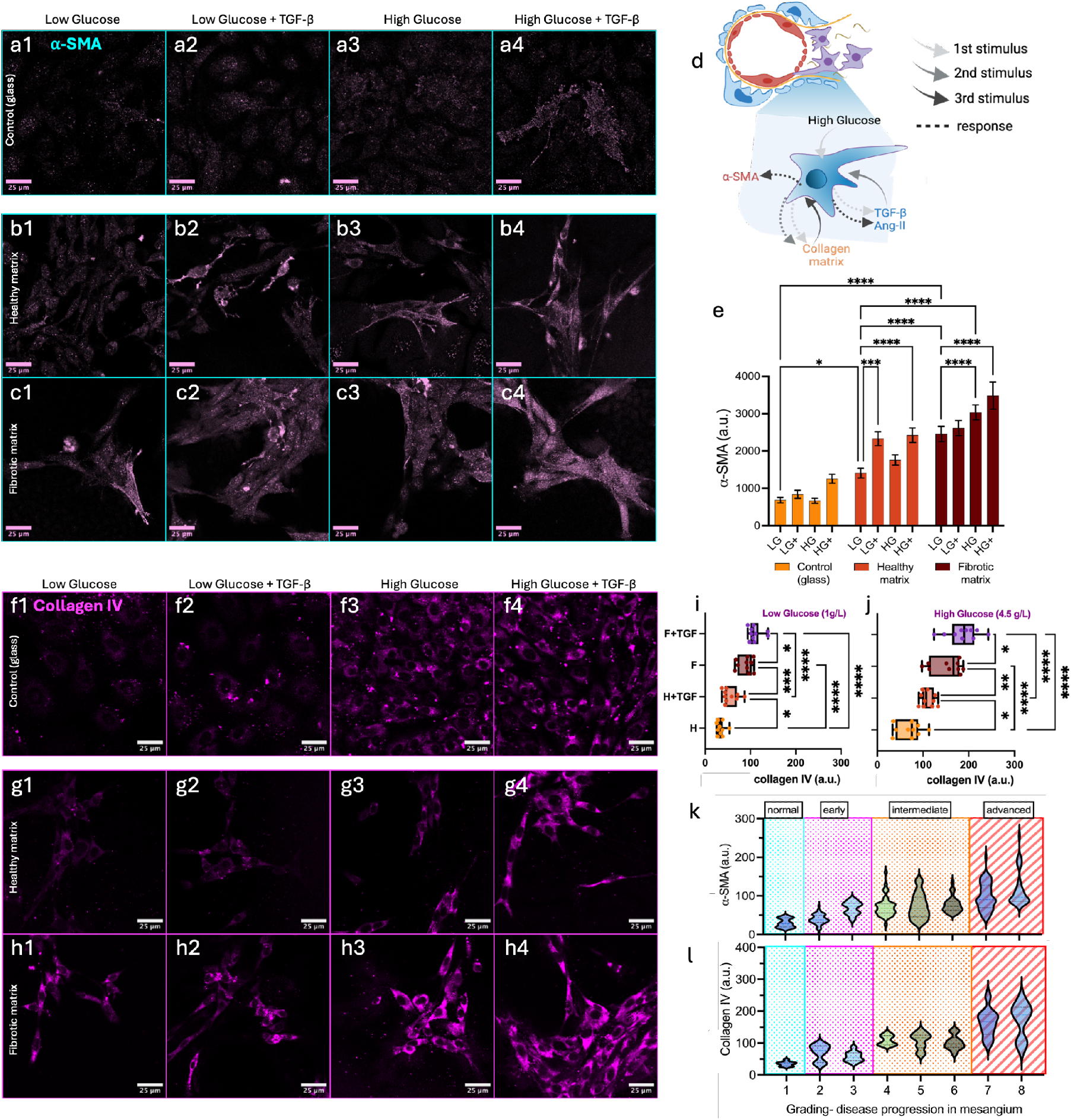
Mechanical stiffness and chemical cues synergistically regulate *α*-SMA and Collagen IV expression in a 3D mesangium model of diabetic kidney disease. (a1–a4) Immunofluorescence images of *α*-SMA expression in cells grown on control (glass) under varying glucose and TGF-*β* 1 conditions. (b1–b4) *α*-SMA expression on soft (healthy-like) matrix. (c1–c4) *α*-SMA expression on stiff (fibrotic-like) matrix. *α*-SMA iexpression increases progressively with both biochemical (high glucose, TGF-*β* 1) and mechanical (stiff matrix) stressors. (d) Schematic diagram shows how matrix stiffness, glucose levels, and inflammation affects DKD progression. It also shows the progression can be assessed by expression levels of *α*-SMA and Collagen. (e) Quantification of *α*-SMA expression shows significant upregulation in the fibrotic matrix, particularly under high glucose and TGF-*β* 1 stimulation. (f1–f4) Collagen IV expression on the control (glass). (g1–g4) Collagen IV on healthy matrix. (h1–h4) Collagen IV on fibrotic matrix. (i–j) Quantification of Collagen IV expression under low (1 g/L) and high (4.5 g/L) glucose confirms mechanical stiffness amplifies TGF-*β* 1-induced ECM deposition. Each data point is a mean of one hundred cells in each of the ten replicates. (k–l) Expression level-based stratification of *α*-SMA and Collagen IV expression. As key DKD biomarkers, they are correlated with disease stages from normal to advanced DKD. Mechanical stiffness contributes significantly to transitions across stages, highlighting its role as a critical driver of disease progression. Statistical test done with 2-way ANOVA, multiple comparisons between each category is done, only significant groups highlighted in the plot. HG=high glucose, LG=low glucose. ns (not significant): 0.123, *: p 0.032, **: 0021, ***: 0.0002, ****: <0.0001.

We booleanized the three mechano-chemical conditions to represent 8 disease states, such as 1 = 000 (no diabetes, no TGF, normal stiffness), 2 = 100 (diabetes only), 3 = 010 (TGF only), 4 = 110 (diabetes + TGF), 5 = 001 (high stiffness only), 6 = 011 (TGF + high stiffness), 7 = 101 (diabetes + high stiffness), 8 = 111 (diabetes + TGF + high stiffness). Mapping the expression of key severity markers *α*-SMA and Collagen IV to disease states (Figure 3k–l, supplementary Table 2 and 3) we capture progressive DKD phenotypes. A notable observation is that the stiffness alone is sufficient to shift cellular behavior toward intermediate and advanced stages, emphasizing its central role in disease progression alongside chemical triggers.

### A combinatorial model reveals synergistic drivers of DKD severity and enables continuous prediction

To enable predictive modeling of fibrotic progression in diabetic kidney disease (DKD), we extended the quantitative analysis of *α*-SMA and Collagen IV severity derived from our 3D in vitro mesangium model (Figure 4). These severity scores were measured across eight defined conditions combining low or high glucose, TGF-*β* 1 presence or absence, and healthy or fibrotic GelMA matrix stiffness. Each data point represents a biological replicate, averaged from 100 cells per condition. Ten replicates per group provided sufficient resolution to capture condition-specific trends while maintaining feasibility for high-content modeling. Though moderate in size, this dataset was amenable to predictive analysis through a probabilistic modeling approach that explicitly accounted for experimental variability and data sparsity. To structure the modeling input, we encoded experimental groups in binary form (0 = low/absent, 1 = high/present) for glucose, TGF-*β* 1, and matrix stiffness (Figure 4a). Severity scores for both markers rose systematically across this parameter space (Figure 4b–g), with grouped and pairwise analyses (Figure 4d,g,h–m) revealing synergistic effects among the three cues. In particular, matrix stiffness amplified the severity of biochemical stress, with *α*-SMA showing the most pronounced synergy. Technical validation of the Gaussian process regression method, including data augmentation strategies is shown in supplementary Figure S10. To build a predictive framework from these sparse measurements, we employed probabilistic model based on Bayesian Gaussian Process (GP) regression. This approach infers a continuous severity surface across the 3D input space without relying on a predefined function. The model was trained using a Markov Chain Monte Carlo (MCMC) sampling scheme with 20,000 iterations, allowing thorough exploration of the parameter space. To prevent overfitting and enhance generalizability given the limited sample size, mild Gaussian noise was added to the input data at each iteration. This data augmentation strategy helped stabilize the model and ensured convergence toward a robust predictive distribution. The final model output (Figure 4n) visualizes the mean predicted severity across all input combinations. The resulting severity surface reflects a smooth, non-linear escalation (Figure 4o) from healthy to pathological states with high glucose acting as a strong global driver and TGF-*β* 1 and matrix stiffness shaping the local topology of the response. Further insight into the GP modeling approach, including posterior probability density distributions of model parameters, is provided in supplementary Figure S11. A 360-degree view of the severity model is presented in the supplementary video. This data-driven framework not only integrates complex mechano-chemical interactions into a single predictive map but also enables interpolation to intermediate or untested conditions (Figure 4p) —offering a versatile tool for stratifying DKD severity and guiding future in vitro or translational studies.

**Figure 4.**
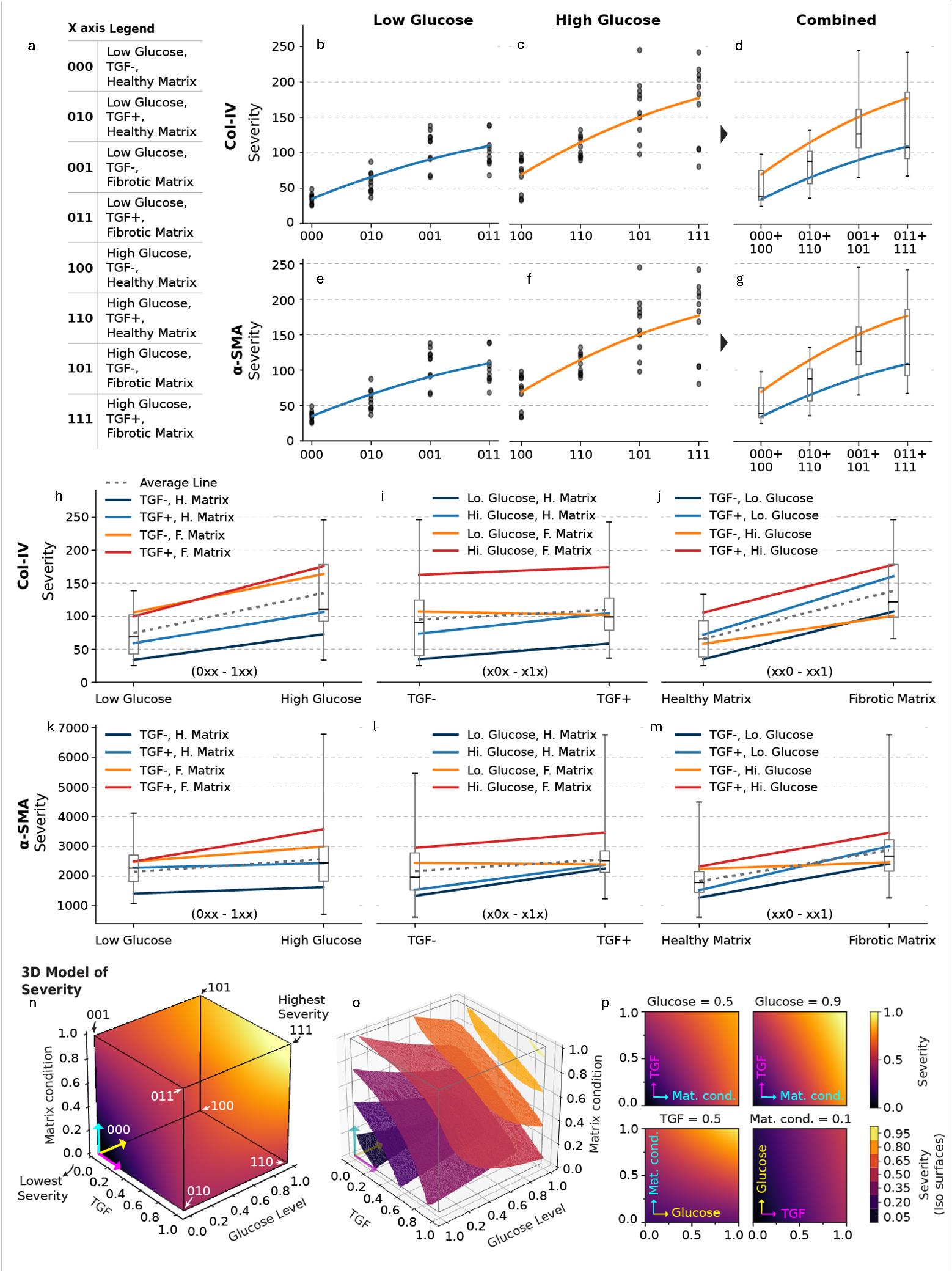
A data-driven severity model stratifies diabetic kidney disease states across eight mechano-chemical conditions and predicts intermediate disease progression. (a) Encoding of eight experimental conditions using binary representation for glucose level (0 = low, 1 = high), TGF-*β* presence (0 = -, 1 = +), and matrix stiffness (0 = healthy, 1 = fibrotic). (b–c) Collagen IV (Col-IV) severity scores across increasing binary codes under low and high glucose, respectively, and (d) grouped plot combining both. (e–f) *α*-SMA severity scores under the same stratification as (b–c), and (g) grouped view. (h–j) Pairwise comparisons showing effects of glucose (h), TGF-*β* (i), and matrix condition (j) on Col-IV severity. (k–m) Equivalent comparisons for *α*-SMA severity. Each line represents the trend across one of the four combinatorial categories. Error bars represent standard deviation. (n) Three-dimensional Gaussian process regression-based model of severity prediction, with mean severity values visualized across the parameter space defined by glucose, TGF-*β*, and matrix condition. (o) Iso-surfaces are shown to display the non-linearity of severity conditions with the combination of the three parameters. (p) 2D images at different severity coordinates show differential impacts of the individual parameters.

## DISCUSSION

Diabetic kidney disease (DKD) continues to be diagnosed far too late, often only after irreversible nephron loss has occurred^26,27^. This delay largely stems from an inability to detect pathophysiological transitions in mesangial behavior—an upstream and mechanochemically active microenvironment critical to glomerular filtration—before structural damage sets in. In this work, we asked whether an engineered 3D in vitro platform can recapitulate these early mesangial manifestations and translate cellular responses to disease-relevant stressors into a predictive diagnostic framework. Our findings support this proposition. We established a high-fidelity 3D hydrogel model of the kidney mesangial microenvironment using GelMA matrices engineered to span the physiological stiffness range of healthy (*∼*0.4–3 kPa) to fibrotic (8-30 kPa) tissue. While the mesangium is small, deeply situated, and experimentally inaccessible in vivo, its strategic location at the vascular–glomerular interface makes it a powerful sentinel of early DKD pathogenesis. This microenvironment is both chemically^28,29^ and mechanically^30,31^ active; mesangial cells both secrete ECM components and respond to mechanical stress, forming a feedback loop that exacerbates disease. Thus, a model capturing both mechanical fidelity and controlled biochemical variation was necessary.

We chose GelMA not only for its tunability but also for its compatibility with mesangial biology. Unlike rigid synthetic scaffolds, GelMA presents integrin-binding RGD motifs and allows for tuning of stiffness, crosslinking density, viscoelasticity, and thermal behavior. Given the known variability in commercial GelMA preparations, we developed our in-house synthesis protocol with controlled methacrylation and characterized our hydrogels extensively. Compression testing, shear rheology, frequency-dependent storage modulus (*G*^*′*^), and differential scanning calorimetry confirmed that 5% and 10% GelMA effectively emulate healthy and fibrotic mesangial stiffness.

Once the mechanical substrate was established, we evaluated mesangial behavior using both label-free and molecular methods. Quantitative phase imaging via GLIM revealed that cells cultured in stiff matrices exhibited nearly 1.7 × higher dry mass accumulation than those in soft gels—indicating enhanced biosynthetic activity, likely reflecting increased ECM deposition. This was confirmed by immunostaining for *α*-SMA and collagen IV, which were significantly upregulated in response to increased stiffness, with or without accompanying high glucose or TGF-*β* exposure.

Importantly, we found that stiffness alone—independent of glucose or inflammatory stimuli—was sufficient to drive fibrotic transitions, but the combination of mechanical and biochemical cues led to synergistic increases in fibrotic marker expression. While elevated glucose and TGF-*β* levels certainly contribute to DKD pathology, their effects were most pronounced when overlaid on a mechanically dysregulated matrix. This suggests that DKD progression may follow a threshold-like mechanism, where once sufficient matrix stiffening occurs, further disease advancement becomes self-reinforcing and less responsive to biochemical correction. This aligns with clinical observations: even when glucose levels are controlled interventionally, fibrosis often continues due to persistent ECM stiffening^32^. Notably, we observed that mesangial cells cultured on control (which are orders of magnitude stiffer than tissue) did not display comparable upregulation of fibrotic markers—highlighting that it is functional, tissue-like stiffness, not absolute stiffness, that governs cellular phenotype.

Beyond fibrotic readouts, we also analyzed mesangial morphology and motility. The cellular morphology also proves to be a valuable tool in understanding how the contractility (*α* − *SMA* expression) affects cell interaction and motility^33^. Under fibrotic and inflammatory conditions, cells lost their characteristic branching^34^ and reticulated morphology, instead forming compact, clustered networks with impaired filopodia extension and reduced directional migration. These features mirror early fibrotic changes seen in vivo and are seldom captured in DKD models, especially not with this level of control and quantifiability. These morphological changes—particularly loss of directional plasticity and increased compactness—may serve as behavior-based biomarkers preceding histological fibrosis, offering a new axis for subclinical disease detection. To move from experimental readouts to a clinically meaningful framework, we constructed a predictive model using Bayesian Gaussian process regression. This approach allowed us to map disease severity as a continuous function of three variables: ECM stiffness, glucose, and TGF-*β* . Each of our eight experimental conditions was encoded as a triplet within this parameter space. Severity scores, derived from *α*-SMA and collagen IV expression, were then used to generate a “severity cube”—a 3D predictive landscape capable of interpolating disease states between our tested conditions. Gaussian process regression was chosen for its flexibility in modeling sparse, nonlinear datasets and its capacity to infer biologically plausible intermediate conditions—a critical advantage given the limited number of feasible in vitro combinations. This model-less regression framework was selected because no explicit mechanistic function could describe the multivariate severity landscape of DKD, making Gaussian processes ideal for capturing complex, organic relationships. We implemented the regression using a radial basis function (RBF) kernel to encode the covariance between data points, capturing smooth nonlinear dependencies. Hyperparameters governing kernel behavior—such as length scale—were optimized using Markov Chain Monte Carlo (MCMC) sampling to locate the highest-likelihood posterior distribution over model functions.

The severity cube provides a novel conceptual and computational bridge between cell biology and clinical diagnostics. It highlights a sharp transition from health to disease along the stiffness axis, with glucose and TGF-*β* acting as accelerants. To further enhance generalizability, we employed Gaussian noise augmentation during model training, reducing the risk of overfitting to limited discrete data points. Specifically, Gaussian noise with variance *σ* ^2^ = 0.1/e was introduced at each MCMC step to mitigate overfitting by encouraging the model to learn robust trends rather than memorizing sparse inputs. This nonparametric learning framework allows for flexible modeling without overfitting—a key strength given the inherent data sparsity in early-stage disease modeling.

Critically, the severity cube supports potential integration with real-world clinical data. Parameters such as renal stiffness (measurable via ultrasound elastography), serum glucose, and circulating cytokines (e.g., TGF-*β* or inflammatory indicators) are already accessible through routine screening^35^. Our model suggests that these could be sufficient to predict DKD severity when mapped onto a biologically grounded in vitro reference, thus reducing reliance on invasive kidney biopsies and enabling earlier, risk-adjusted therapeutic decisions.

Furthermore, the 3D in-vitro mesangial model and the severity cube is inherently longitudinal: by tracking severity zone shifts over time, it may be possible to monitor disease progression or therapeutic response. This opens new possibilities for patient stratification and real-time adjustment of treatment protocols^36^. Eventually, incorporation of additional biomarkers, such as ECM remodeling enzymes or metabolic activity signatures, could enhance both sensitivity and specificity. Given the modularity of the platform, this approach could also be adapted to other mechanochemically governed diseases such as liver fibrosis, oral mucosal fibrosis, or even cancer progression where ECM remodeling plays a central role.

Nevertheless, some gaps remain. While our 3D model captures key mesangial behaviors with high fidelity, it lacks vascular, immune, and hormonal components known to influence DKD evolution. Future versions of the platform will need to integrate endothelial co-cultures, immune modulators, and potentially fluid flow via microfluidics to approximate in vivo conditions more closely. Moreover, validation with primary mesangial cells from both healthy donors and DKD patients will be essential for clinical translation. Expanding the statistical training set to include a wider range of molecular and phenotypic readouts will enhance the robustness of severity zone classification and model generalizability across populations and ethnicities.

In conclusion, this work demonstrates that early-stage DKD transitions—driven by a triad of mechanochemical stressors—can be faithfully captured in vitro and translated into a predictive diagnostic continuum. This may augment existing therapeutic efficiency and standard practices focused strongly on chemical biomarkers of DKD^37^. Our integrative approach bridges experimental rigor with computational inference, providing a scalable foundation for non-invasive DKD screening and real-time disease monitoring. This approach demonstrates the potential of engineered microenvironments in advancing precision medicine and illustrates the importance of integrating cell biology, biophysics, and statistical learning to better understand disease progression.

## CONCLUSION

This study establishes a predictive, mechanochemically aware in vitro platform that captures early fibrotic transitions in diabetic kidney disease (DKD) through a tunable 3D mesangial model. By integrating matrix stiffness, glucose, and TGF-*β* 1 into a Gaussian process regression framework, we generate a severity cube that maps disease progression across multivariate conditions. This approach bridges experimental modeling and statistical inference to enable non-invasive, quantitative stratification of early DKD. While the model excludes immune, vascular, and hormonal inputs, its modularity allows future expansion with co-cultures and fluidic systems. The platform supports clinical translation through validation with patient-derived mesangial cells and longitudinal correlation with biomarkers. In summary, this work offers a scalable foundation for predictive diagnostics in DKD and presents a generalizable strategy for modeling fibrotic diseases influenced by mechanical cues.

## MATERIALS AND METHODS

### Materials

The antibodies and chemicals used in this study were rabbit polyclonal anti-*α* smooth muscle actin from Abcam; rabbit anti-mouse polyclonal anti-Collagen-IV from MD Biosciences; goat anti-rabbit with AlexaFluor 488 and 647 from Invitrogen; diamidino-2-phenylindole (DAPI), phalloidin–Alexa 647, goat serum, and TGF-*β* from ThermoFisher Scientific; phalloidin–Atto 565, gelatin Type A (bloom 300), methacrylate anhydride (MAA), lithium phenyl-2,4,6-trimethylbenzoylphosphinate (LAP), sodium hydroxide (98%), hydrochloric acid (HCl), paraformaldehyde (prilled, 95%), Tween-20, and commercial GelMA (80% and 40%), ninhydrin from Merck. Dulbecco’s Modified Eagle Medium (DMEM)–high glucose (4.5 g/L), DMEM–low glucose (1 g/L), and fetal bovine serum (FBS)–heat-inactivated, antibiotic–penicillin–streptomycin, ethanol, and methanol were purchased from Avantor. Human renal mesangial cells-healthy primary cell line (P10661, Innoprot) passage 3-5 and Mouse kidney mesangial cell line DPK-KMGC-M (passage 8 to 15) were used for cell culture and were all commercially acquired. For compatibility testing mouse mesangial cells and human primary cells were used, and for model prediction data, human primary cell results were used.

### GelMA Synthesis

Gelatin methacryloyl (GelMA) was synthesized in PBS with pH 7.4 at 10% w/v. Gelatin type A bloom 300 was dissolved in PBS and heated at 50 °C using a hotplate and stirred continuously using a magnetic stirrer. The gelatin solutions were adjusted to *pH*9 by the addition of 5*N* sodium hydroxide (NaOH) to promote the reaction of the amino groups on the gelatin. Methacrylate anhydride was added to the gelatin solutions at a feed ratio of 0.1 : 1 (ml of MAA to g of gelatin) at the rate of 0.5*ml/min* while stirring at 50 °C. The addition of MAA was divided into 6 separate times with an average of 10 min between each addition. As the reaction proceeded, the pH decreased. As such, the pH of the reaction solutions was adjusted to pH 9 after each addition of MAA for optimal reaction of gelatin with MAA. After around 1.5*h* of the reaction, 1*N* HCl was added to the solutions to be adjusted to the physiological pH of 7.4. All the reactions were carried out in the dark. The solutions were then filtered with a 70 mm paper filter, followed by an antibacterial filter (0.2*µm*). Then, the solutions were dialyzed for about 4 days against Milli-Q water at 40 °C using a dialysis membrane (12 kD molecular weight cut-off). This was done to filter out small molecules of less than 12 kDa, unreacted MAA, salts, and methacrylic acid byproducts. After dialysis, the solutions were placed at *−*70 °C for 3 days through freeze-drying and finally lyophilized to obtain the dry GelMA products.

### Hydrogel preparation and characterization

50*mg/ml* LAP photoinitiator was prepared by dissolving in sterile PBS pre-warmed to 60 °C. To homogenously dissolve the LAP the mixture was pulse ultrasonicated for 5 − 10 minutes and then maintained at 60 °C in a water bath. 5% and 10% (w/v) GelMA hydrogels were dissolved in sterile PBS pre-warmed to 60 °C. The GelMA precursor solution was made by adding the photoinitiator LAP such that the final concentration was 5*mg/ml* (or 0.5%*w/v*). The precursor was gently mixed by pipetting avoiding the bubbles from forming. The precursor solution was kept at 37 °C before UV crosslinking to prevent it from becoming viscous and for easy spreadability. 130*µl* of the precursor solution was dropped in the center of the 35mm glass bottom dish within the 20mm center well of the dish. A pipette tip was used to ensure that the solution was evenly distributed up to the walls of the well. To crosslink the GelMA, a UV LED lamp with a central wavelength of 365*nm* was used to expose the dishes containing the precursor solution for 50 seconds. 1-2 ml pre-warmed PBS at 37 °C was added to the crosslinked GelMA and kept in the incubator for 10 − 15 min. The PBS was aspirated to remove all non-crosslinked GelMA and wash out initiators.

### 3D cell culture

To create healthy and DKD mesangium in 3D hydrogel culture, 5% and 10% GelMA were used to mimic healthy and fibrotic mesangium, respectively. These match the elastic modulus of the healthy and pathological conditions found in the literature^14,15^. The GelMA is entirely protein-based and has RGD motifs that ensure cell adhesion like that in the tissues^38^. Mesangial cells were cultured in low glucose and high glucose conditions for at least two passages before being added to dishes under non-diabetic control and diabetic conditions. Cells were trypsinized and sufficiently neutralized with fresh media. After the GelMA was washed in warm PBS, 10000 mesangial cells were added only to the central well of the hydrogel dishes to achieve 20 − 30% confluence. No more than 100*µl* was added to each well to avoid spilling cells onto other parts of the dishes. This ensured that there were no effects from cells growing on the non-hydrogel areas of the dishes. The cells were allowed to adhere to the hydrogel for 2 hours, after which fresh pre-warmed low or high glucose media (1–2 ml) was added to the respective dishes. The cells were allowed to spread overnight, and the old media was then replaced with fresh media. The total study duration was 5 days (120 hours). TGF-*β* 1 was added at a concentration of 20*ng/ml* to the dishes with inflammatory-type conditions on the first day. For the next 4 days, the cells were regularly supplemented with 10*ng/ml* of TGF-*β* 1 to simulate a consistent level. On the 5_*t*_*h* day, live imaging of the cells was performed to measure the effect on mesangial cell processes and their cell-cell and cell-matrix interactions.

### Quantification of degree of substitution

Fourier Transform Infrared (FTIR) spectra were collected using a PerkinElmer ATR spectrophotometer (4000–400 cm^*−*1^, 16 scans, 2*cm*^*−*1^ resolution) at room temperature. Characteristic peaks confirmed methacrylation of gelatin, with intensity differences between 5% and 10% GelMA indicating variations in crosslink density. The degree of substitution (DoS) for our GelMA hydrogels was 95% using the Ninhydrin assay^39^. The absorbance was measured using a UV-visible spectrometer at 570nm wavelength. We used commercial-grade GelMA with 40% and 80% DoS (Sigma-Aldrich), along with blanks (no protein) and a positive control (only gelatin, no substitution), to create a standard curve. The GelMA absorbance was determined by fitting the absorbance value to the standard curve and obtaining the corresponding DoS. To confirm, Proton Nuclear Magnetic Resonance (^1^H NMR) spectra were recorded on a 400 MHz Bruker Avance instrument. Dried GelMA powder was dissolved in D_2_O (1% w/v). The analysis confirmed methacrylation by identifying characteristic peaks from methacrylate groups and the gelatin backbone, used to estimate the degree of substitution.

### Mechanical characterization of mesangium relevant hydrogels

Rheological measurements of hydrogels were performed on an Anton Paar MCR 102 rheometer using parallel plate geometry. Temperature sweeps were conducted at 1 Hz from 5–40 ^*°*^C under 0.1 N normal force. Frequency sweeps were performed at room temperature with 0.1% shear strain over 0.01-100 Hz. Compression testing of GelMA hydrogels was performed using a Mark-10 Plug and Test™ system equipped with an FS05-100 force sensor. Unconfined compression tests were conducted at two different strain rates: 5 mm/min and 1 mm/min. The hydrogels were compressed until slightly beyond their breaking point under a constant crosshead speed. This approach was intended to assess the viscoelastic response and mechanical integrity of the hydrogels under both physiologic and impact-like loading conditions. Differential Scanning Calorimetry (DSC) was performed using a NETZSCH DSC 200 F3 under nitrogen (100 mL/min) with a heating rate of 10 ^*°*^C/min. Samples were sealed in standard aluminum pans. Endothermic transitions were used to assess water loss, protein denaturation, and thermal stability differences between 5% and 10% GelMA hydrogels.

### Gradient Light Interference Microscopy and drymass analysis

Quantitative phase imaging was performed using GLIM mode integrated into a Nikon Ti2-E inverted microscope (Phi Optics, USA) equipped with a 40×/0.95 NA air objective lens. Phase-shifted images were captured using a Hamamatsu ORCA-Fusion sCMOS camera with a native pixel size of 0.16 *µ*m, enabling high-resolution label-free phase contrast imaging. All images were acquired in transmission mode under standard cell culture conditions. The acquired raw GLIM images were phase-unwrapped and reconstructed by the GLIM acquisition software (Phi Optics) to yield quantitative phase maps in radians. Subsequent processing and dry mass analysis^19,40^ were conducted in MATLAB R2024b.

#### Optical Path Length (OPL) Calculation

The unwrapped phase shift *φ* (*x, y*) at each pixel was converted into optical path length (OPL) using the formula:

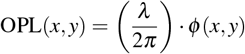

where *λ* = 550 nm is the illumination wavelength. OPL values represent the integrated refractive index contrast between the sample and the surrounding medium.

#### Dry Mass Computation

Cellular dry mass was computed from the OPL maps using the relationship between refractive index and biomass content, based on the Gladstone-Dale equation^19^:

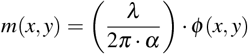

where *m*(*x, y*) is the dry mass density (in pg/*µ*m^2^) and *α* = 0.18 *µ*m^3^/pg is the refractive increment for cellular proteins. This yields a dry mass map in units of pg/*µ*m^2^, integrated over the cell thickness.

To account for variations in the optical path introduced by the substrate or surrounding gel matrix, background-normalized dry mass maps were calculated. For each image, a background region of interest (ROI) was manually selected in the adjacent hydrogel devoid of cells. The mean background dry mass was computed and subtracted from the raw dry mass map. This normalization highlights true biomass accumulation attributable to cellular presence and excludes contributions from hydrogel or surface artifacts.

### DIC Imaging, Filopodia Analysis, and Directional Plasticity Quantification

Label-free differential interference contrast (DIC) imaging was performed using a GE DeltaVision inverted microscope equipped with a 10×/0.40 NA air objective. Mesangial cells were imaged live at 37 ^*°*^C under standard culture conditions to capture morphological dynamics in response to matrix stiffness and biochemical stimuli. High-resolution DIC images were acquired at multiple time points to assess static morphology and real-time filopodia dynamics^41^. All image processing and quantitative analysis were performed in MATLAB R2024b and FIJI (ImageJ). Images were first background-corrected using a rolling ball filter and contrast-normalized using linear histogram stretching. For each experimental condition, at least 100 cells were analyzed across five independent biological replicates.

#### Filopodia Length Analysis

Filopodia were detected by applying a directional edge-enhancement filter followed by morphological skeletonization. Lengths were computed by tracing individual filopodial processes extending from the cell body. The centroid of each cell was used as a seed, and filopodia were segmented based on curvature and local intensity gradients. The final filopodial length for a cell was computed as:

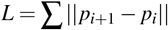

where *p*_*i*_ and *p*_*i*+1_ are adjacent coordinates along the skeletonized filopodium.

#### Compactness Measurement

To quantify the spatial distribution of mesangial cells, compactness was assessed by analyzing local cell density and gap fraction within a defined region. For each field of view, a 200 × 200 *µ*m^2^ ROI was selected in the central part of the image. Binary masks of cell-covered regions were generated using adaptive thresholding on the DIC image followed by morphological opening to remove background noise. Compactness (%) was calculated based on the inverse of the background (gap) area within the ROI:

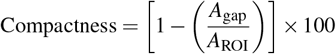

where *A*_gap_ is the total area not covered by cells, and *A*_ROI_ is the area of the 200 × 200 *µ*m^2^ window. A value close to 100% indicates high compactness (dense, gap-free clustering), while lower values reflect sparse cell distribution with visible intercellular gaps.

#### Filopodia Directionality and Angular Spread

To quantify the directional plasticity of filopodia, time-lapse DIC images were acquired over 24 minutes. For each frame, the angular orientation of filopodial projections was computed with respect to the cell centroid using vector decomposition. The angular spread (Δ*θ*) for each cell was defined as the standard deviation of projection angles:

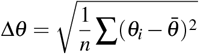

where *θ*_*i*_ is the angle of the *i*^th^ filopodium, and 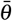 is the mean orientation. Lower angular spread indicates directional bias (e.g., compact, bundled protrusions), while higher spread reflects exploratory, multidirectional filopodial dynamics. This metric was used to compare mesangial cells across healthy and fibrotic conditions. All measurements were averaged across 5–10 frames per time-lapse movie, and statistical comparisons were made using ANOVA with Bonferroni correction for multiple conditions.

### Cell Fixation and Immunolabeling

Cells in the hydrogel matrix were fixed in chilled methanol. 100% methanol was pre-chilled in a -20 °C freezer for 30 minutes. The media was aspirated from the dishes, and the samples were washed 2–3 times with PBS. After removing all PBS, 1 ml of chilled methanol was added to each dish, ensuring complete coverage of the hydrogel. The dishes were then placed in the -20 °C freezer for 10 minutes to allow fixation. Following fixation, the cells were washed and stored in a 4°C refrigerator in PBS. For immunolabeling, the cells were permeabilized with 0.2% Triton X-100 in PBS for 10 minutes, followed by gentle washing three times with PBS to avoid injury or rupture of the hydrogel. The cells were blocked with 5% goat serum in PBS for 35-45 minutes at room temperature. The primary and secondary antibodies were diluted as recommended by the manufacturer in 5% goat serum in PBS. The primary antibody was added to the hydrogel wells (1:400 dilution for *α*-SMA and Collagen-IV) and incubated in the dark for 2 hours at room temperature followed by three PBS washes. The secondary antibody was then incubated in the dark for 1 hour at room temperature followed by three PBS wash. The secondary antibodies was added at 1:500 dilutions and kept at room temperature for an hour. DAPI was used to stain the nucleus for 15 minutes. The cells were finally washed in PBS and stored in a 4°C fridge in the dark condition before imaging. Confocal fluorescence imaging was performed within a week of labeling.

#### Confocal Imaging and Analysis

Confocal laser scanning microscopy was used to quantify protein expression levels of *α*-smooth muscle actin (*α*-SMA) and Collagen IV in mesangial cells embedded in 3D hydrogels. Imaging was performed on a Zeiss LSM 800 confocal microscope equipped with a 40×/1.20 NA water immersion objective. The pinhole was set to 1 Airy unit, and excitation was achieved using a 0.5% laser power setting to minimize photobleaching while preserving signal-to-noise ratio. To ensure quantitative comparability, all imaging parameters and labeling conditions were kept constant across experimental groups. Only surface-adhered cells (top ∼30–40 *µ*m of the 240 *µ*m hydrogel) were imaged to maintain uniform optical thickness across conditions and avoid depth-dependent signal attenuation. For each condition, *N* = 10 samples were analyzed, consisting of 5 biological replicates and 5 technical replicates. Within each replicate about 10 images were taken from different regions randomly. From these images we selected 100 randomly chosen cells for quantification, and the mean fluorescence intensity within each was quantified. Background subtraction was performed using local rolling ball correction, and ROI fluorescence was normalized to background-corrected intensities. Fluorescence intensity values for *α*-SMA and Collagen IV were pooled across replicates for statistical comparison. Cells were excluded if overlapping nuclei or partial visibility compromised the ROI quality.

### Statistical Modeling and Gaussian Process Regression

To quantitatively predict fibrotic severity across multivariate mechano-chemical conditions in our 3D mesangium model, we implemented a Bayesian Gaussian Process Regression (GPR) framework. This non-parametric approach was selected due to its ability to model complex, nonlinear interactions without assuming an explicit functional form, making it especially suitable for sparse datasets derived from physiologically grounded experimental systems.

#### Data Encoding and Preprocessing

Experimental conditions were encoded as binary triplets based on the presence or absence of three key variables: glucose level (0 = low, 1 = high), TGF-*β* 1 stimulation (0 = absent, 1 = present), and matrix stiffness (0 = soft/healthy-like, 1 = stiff/fibrotic-like). Each triplet served as an input vector *x*_*i*_ *∈* ℝ^3^, with the corresponding output *y*_*i*_ representing a severity score derived from the expression intensities of either *α*-SMA or Collagen IV, averaged across 100 cells per biological replicate. For each condition, *N* = 10 samples were used. Within each replicate, regions of interest (ROIs) were selected from 100 randomly chosen cells.

#### Gaussian Process Regression

We modeled the severity function as a stochastic process:

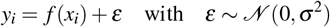

Assuming a Gaussian prior over functions, GPR estimates the posterior distribution of the severity function given the observed data. The kernel (covariance) function determines the smoothness and structure of the predictions. We used the Radial Basis Function (RBF) kernel:

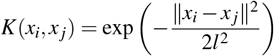

where *l* is the length scale hyperparameter controlling the correlation decay between nearby input points.

#### Hyperparameter Optimization via MCMC

To determine the optimal kernel hyperparameters, we used a Markov Chain Monte Carlo (MCMC) approach with 20,000 iterations per condition. The log-likelihood function was used to guide sampling:

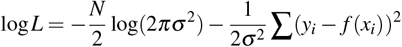

For each MCMC step, a new set of hyperparameters {*l, σ* ^2^ } was sampled, and the likelihood of the observed data under the corresponding GP model was computed. To mitigate overfitting and enhance generalizability given the limited sample size, we implemented data augmentation by adding Gaussian noise with variance:

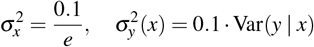

at each iteration, following a scheme designed to broaden the posterior distribution and stabilize convergence.

The final model prediction was computed by averaging the top 50% of MCMC samples with the highest log-likelihood scores, yielding a smoothed severity function capable of interpolating intermediate disease states across the full parameter space.

#### Severity Cube Construction

The trained GPR model was used to generate a 3D severity cube mapping predicted fibrotic severity across the glucose–TGF-*β* –stiffness space. Each axis of the cube corresponds to one of the encoded variables, and the scalar field represents the mean predicted severity at each voxel. This model allows for continuous stratification of disease progression beyond the eight discrete experimental conditions and reveals nonlinear synergistic effects between chemical and mechanical inputs.

## SUPPLEMENTARY INFORMATION

The supplementary information is provided in the supplementary file and videos.

## ACKNOWLEDGEMENTS

The research is funded by the H2020 FET-Open RIA project “OrganVision” 964800. We also acknowledge UiT The Arctic University of Norway for funding the article processing charges of the published article.

## AUTHOR CONTRIBUTIONS

B.G. conceptualized the study, wrote the paper, and performed the experiments. B.G. analysed imaging data. F.N. and B.G. developed and implemented the Baysian model framework. N.K performed the mechanical and chemical characterization of the mesangium relevant hydrogels. K.A.F. contributed to study design and review. K.A. contributed in providing infrastructure and funding. All authors reviewed and revised the manuscript.

## DECLARATION OF INTERESTS

The authors declare no competing interests.

## DATA AVAILABILITY STATEMENT

Data available on request from the authors.

## Notes

### Competing Interest Statement

The authors have declared no competing interest.

### Summary of Updates

In this revision, we have added a mathematical modeling framework that integrates the experimental data and provides a predictive mechanism for determining the severity of the disease condition. This is reflected in Figure 4. We have also introduced a detailed mechanical characterization of the hydrogel substrate material. This is reflected in Figure 1. The earlier figures are now consolidated in Figures 2 and 3. The supplemental files are updated.

