## Supplementary File for "Predicting Early Diabetic Kidney Disease Using a 3D Mesangial Model and Bayesian Mechano-Chemical Stratification"

May 6, 2025

#### Supplementary Materials

#### Supplementary Figures

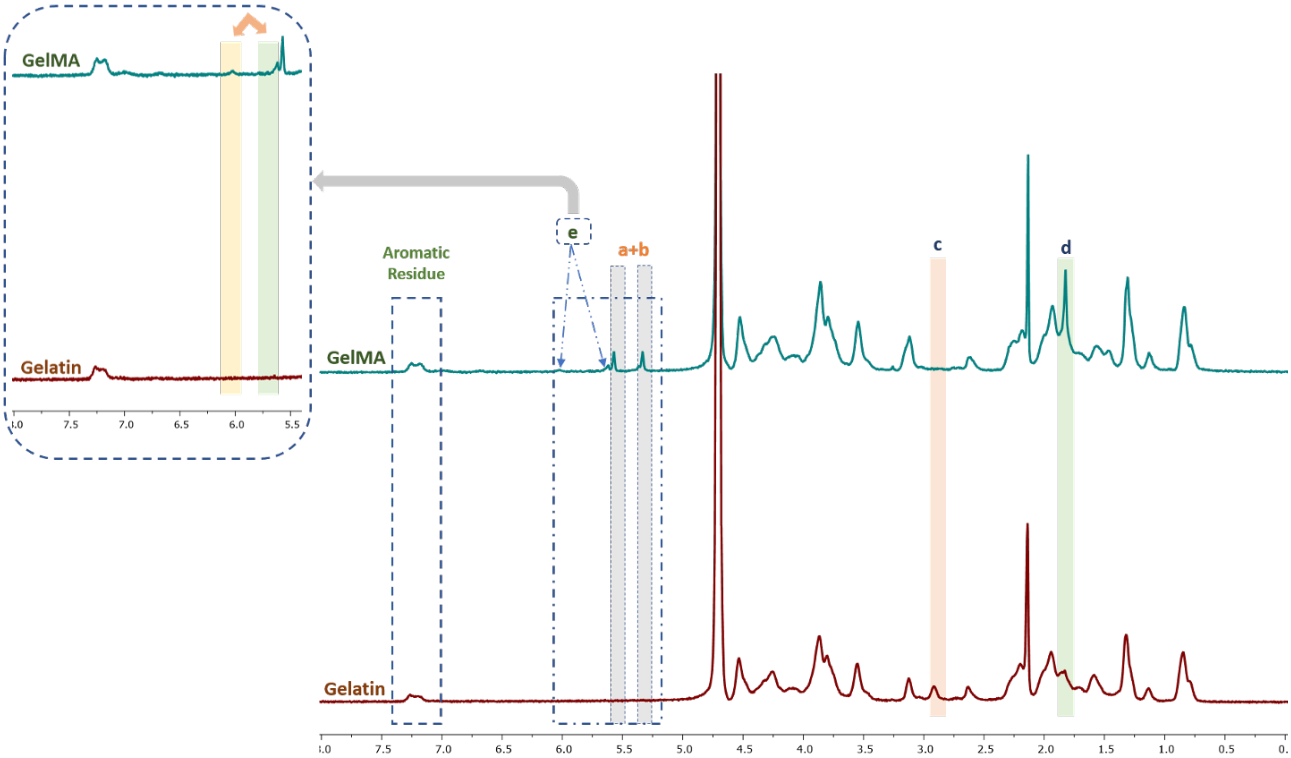


Fig. S1: **$^1$H NMR spectra of Gelatin and GelMA confirming methacrylation.** Comparison of $^1$H NMR spectra of unmodified gelatin (red) and methacrylated gelatin (GelMA, blue-green) shows key spectral differences indicative of successful GelMA synthesis. Peaks labeled **a** and **b** correspond to the methacrylate vinyl protons (5.3–5.7 ppm) and are absent in gelatin, confirming methacrylate functional group addition. Peaks **c** and **d** (1.8–2.0 ppm) arise from the methylene protons adjacent to the methacrylate group. Peak **e** near 7.2 ppm corresponds to aromatic residues present in both gelatin and GelMA. The appearance of new peaks in GelMA spectra and absence in gelatin validates the introduction of methacrylate groups onto gelatin's backbone.

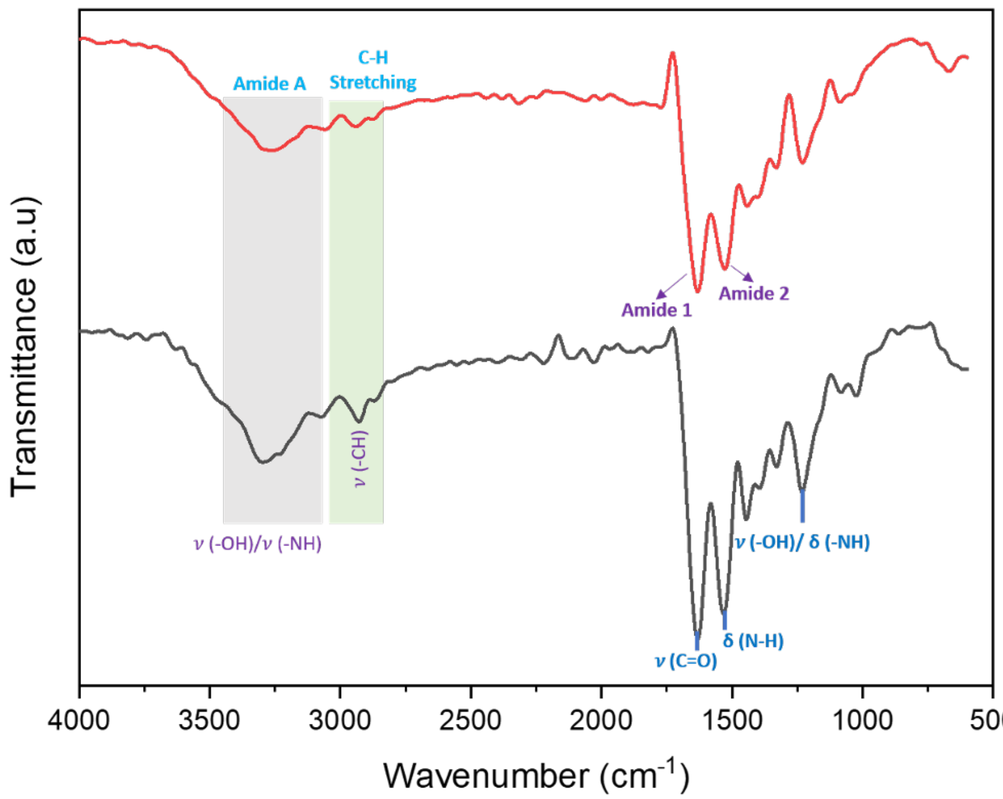


Fig. S2: **Fourier Transform Infrared (FTIR) Spectra. FTIR spectra confirming the successful methacrylation of gelatin..** Representative FTIR spectra of native gelatin (black) and methacrylated gelatin (GelMA, red) highlight characteristic vibrational peaks and chemical modifications upon methacrylation. The broad band in the 3200–3400cm$^{-1}$ region corresponds to stretching vibrations of –OH and –NH groups (Amide A). Peaks around 2900cm$^{-1}$ indicate symmetric and asymmetric C–H stretching. The fingerprint region (1600–1700cm$^{-1}$) shows distinct Amide I and Amide II peaks, associated with C=O stretching and N–H bending. Additional peaks near 1640cm$^{-1}$ and 1530cm$^{-1}$ confirm methacryloyl substitution, consistent with successful GelMA synthesis. The reduction in –OH/–NH absorption in GelMA relative to gelatin indicates methacrylate-induced consumption of reactive amine and hydroxyl groups. Characteristic peaks for C=C and amide bonds are observed, with differences in peak intensities reflecting variations in gel concentration and chemical modification that underpin the mechanical distinctions between 5% and 10% formulations.

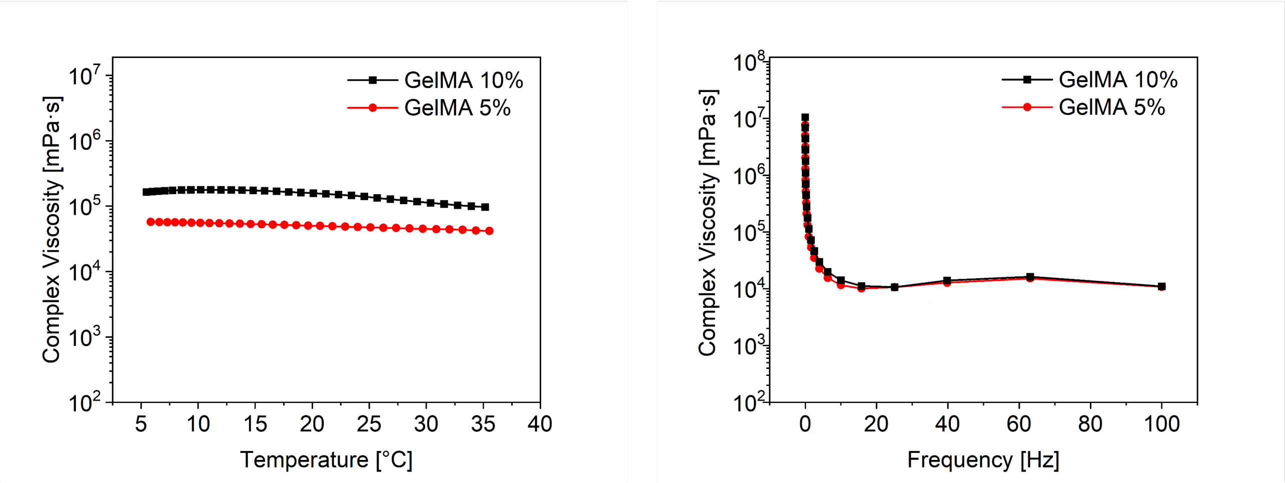


Fig. S3: **Rheological characterization of 5% and 10% GelMA hydrogels**. **(Left)** Complex viscosity of 5% (red circles) and 10% (black squares) GelMA solutions as a function of temperature (5°C to 35°C), measured under constant shear. The 10% GelMA displays consistently higher viscosity, with only a minor reduction at elevated temperatures, indicating greater thermal stability. **(Right)** Frequency-dependent complex viscosity profiles showing pronounced shear-thinning behavior at low frequencies for both GelMA concentrations. At higher frequencies, the viscosity stabilizes, demonstrating frequency-independent viscoelastic properties. These results confirm that 5% GelMA ( 0.4–3kPa) and 10% GelMA ( 8–35kPa) form robust precursors suitable for emulating the stiffness range of healthy and fibrotic kidney mesangial matrices, respectively.

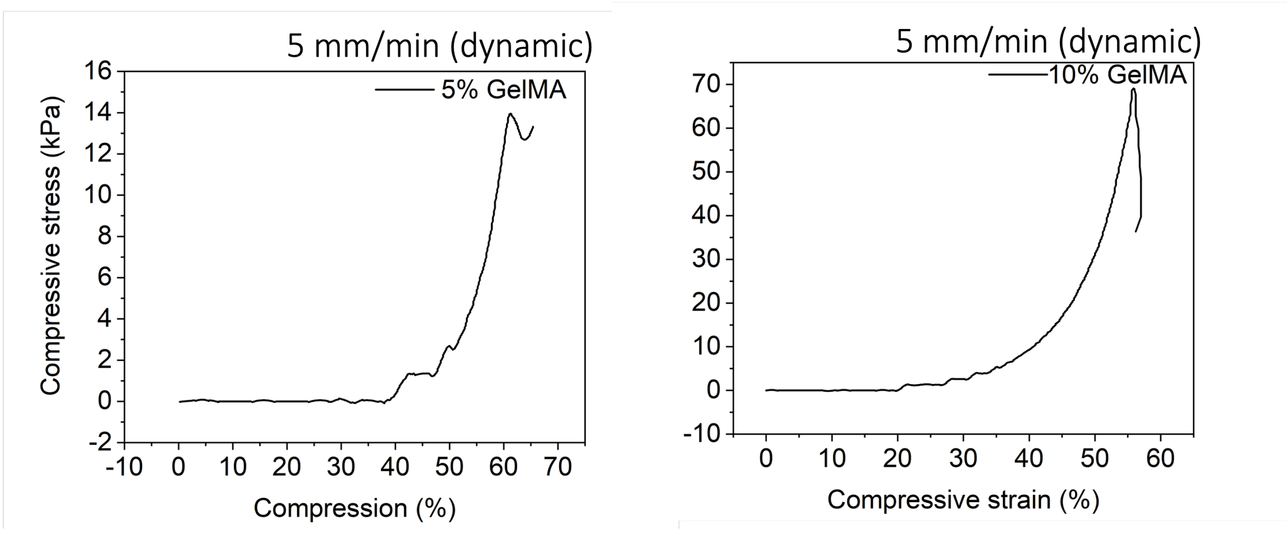


Fig. S4: **Compressive mechanical testing of 5% and 10% GelMA hydrogels in dynamic loading conditions.**. Representative stress–strain curves of 5% GelMA (left) and 10% GelMA (right) hydrogels obtained under dynamic compression at a rate of 5 mm/min. The hydrogels exhibit typical nonlinear behavior, with a long low-stiffness region followed by a sharp increase in compressive stress beyond 50% strain. The 10% GelMA shows a significantly higher compressive modulus and peak stress compared to the 5% GelMA, indicating a stiffer and more mechanically resistant network structure, consistent with increased polymer concentration and crosslink density.

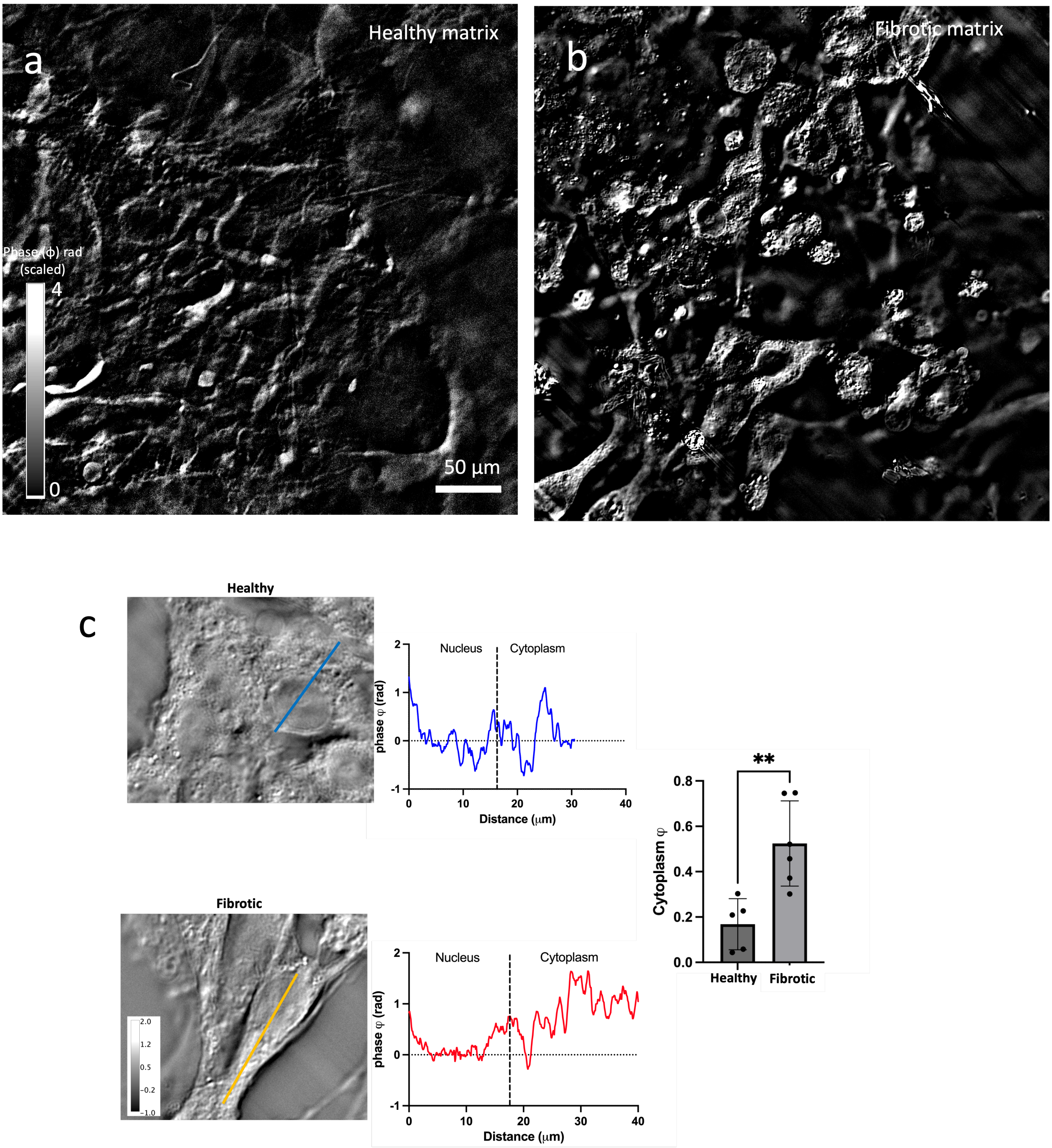


Fig. S5: **GLIM imaging reveals increased phase shift in mesangial cells grown on fibrotic matrix.**. Gradient Light Interference Microscopy (GLIM) images of mesangial cells cultured on (a) 5% GelMA (healthy matrix) and (b) 10% GelMA (fibrotic matrix). The fibrotic matrix condition shows a marked increase in phase shift values, indicative of enhanced dry mass and biosynthetic activity in cells exposed to stiffer extracellular environments. (c) Representative single-cell phase profiles from healthy (top) and fibrotic (bottom) conditions demonstrate a higher cytoplasmic phase signal in fibrotic matrix. Quantification of cytoplasmic phase shift across conditions (right panel) reveals a significant increase in mean cytoplasmic phase in cells on fibrotic matrix compared to healthy (*p $<$ 0.01, unpaired t-test). These results suggest elevated biosynthetic and intracellular density in response to fibrotic mechanical cues.

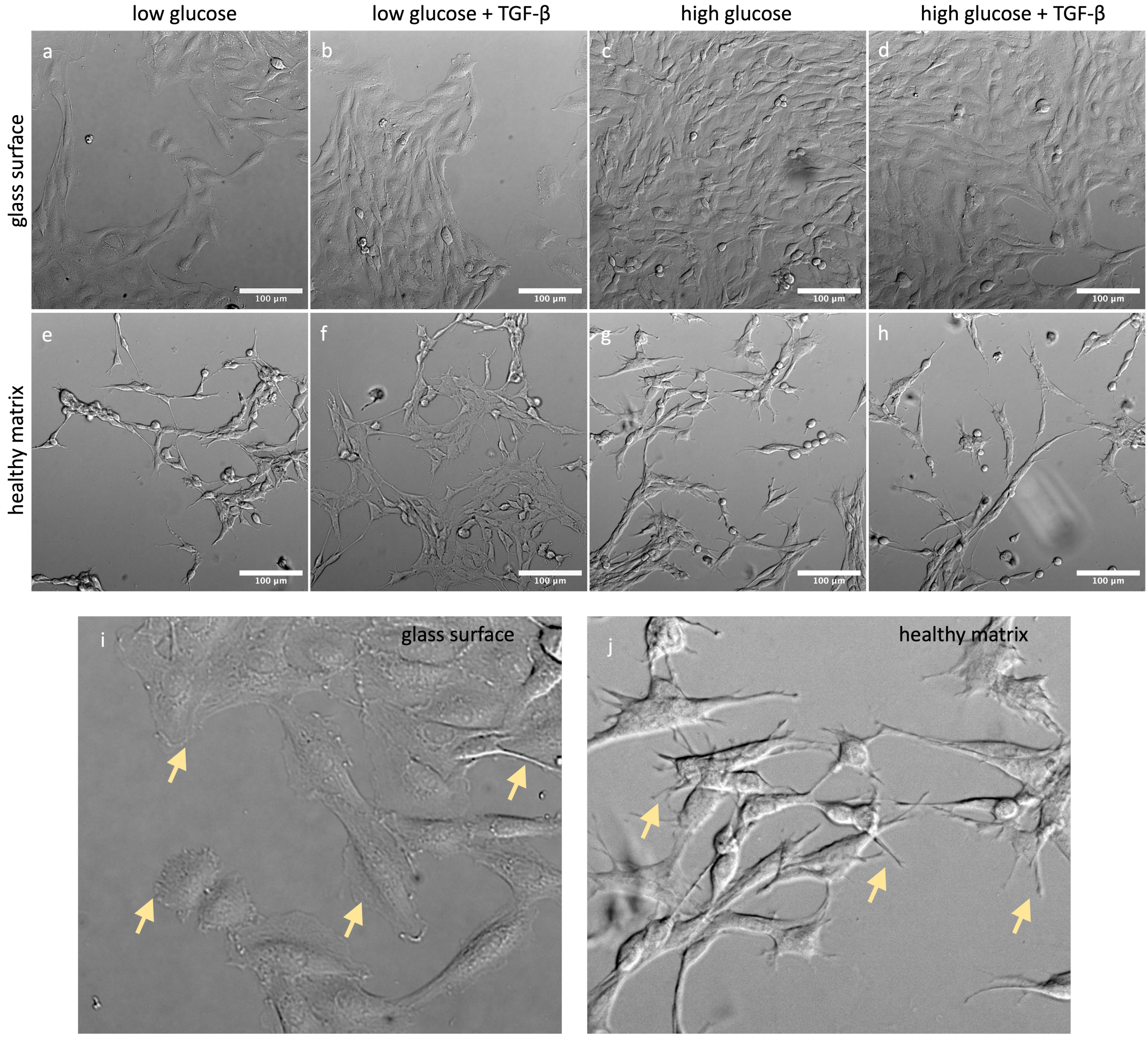


Fig. S6: **Comparison of mesangial cell morphology on 2D glass substrate and 3D GelMA hydrogel**. Representative DIC images of kidney mesangial cells cultured on standard glass surface (a–d) and 5% GelMA hydrogel matrix mimicking healthy mesangial stiffness (e–h), under four chemical conditions: low glucose (a, e), low glucose + TGF-$\beta$ (b, f), high glucose (c, g), and high glucose + TGF-$\beta$ (d, h). Cells on the glass surface display faster proliferation and lamellar morphology, especially under high glucose (c). In contrast, cells on the GelMA matrix exhibit reduced proliferation and retain branched, process-rich morphologies characteristic of mesangial cells. Panels i and j are magnified views showing lamellar processes on glass (i) and branched-elongated mesangial processes in hydrogel matrix (j), marked by yellow arrows. Notably, cell morphology shifts with chemical stimuli even within the 3D matrix, despite the suppression of proliferation under fibrotic-like chemical cues.

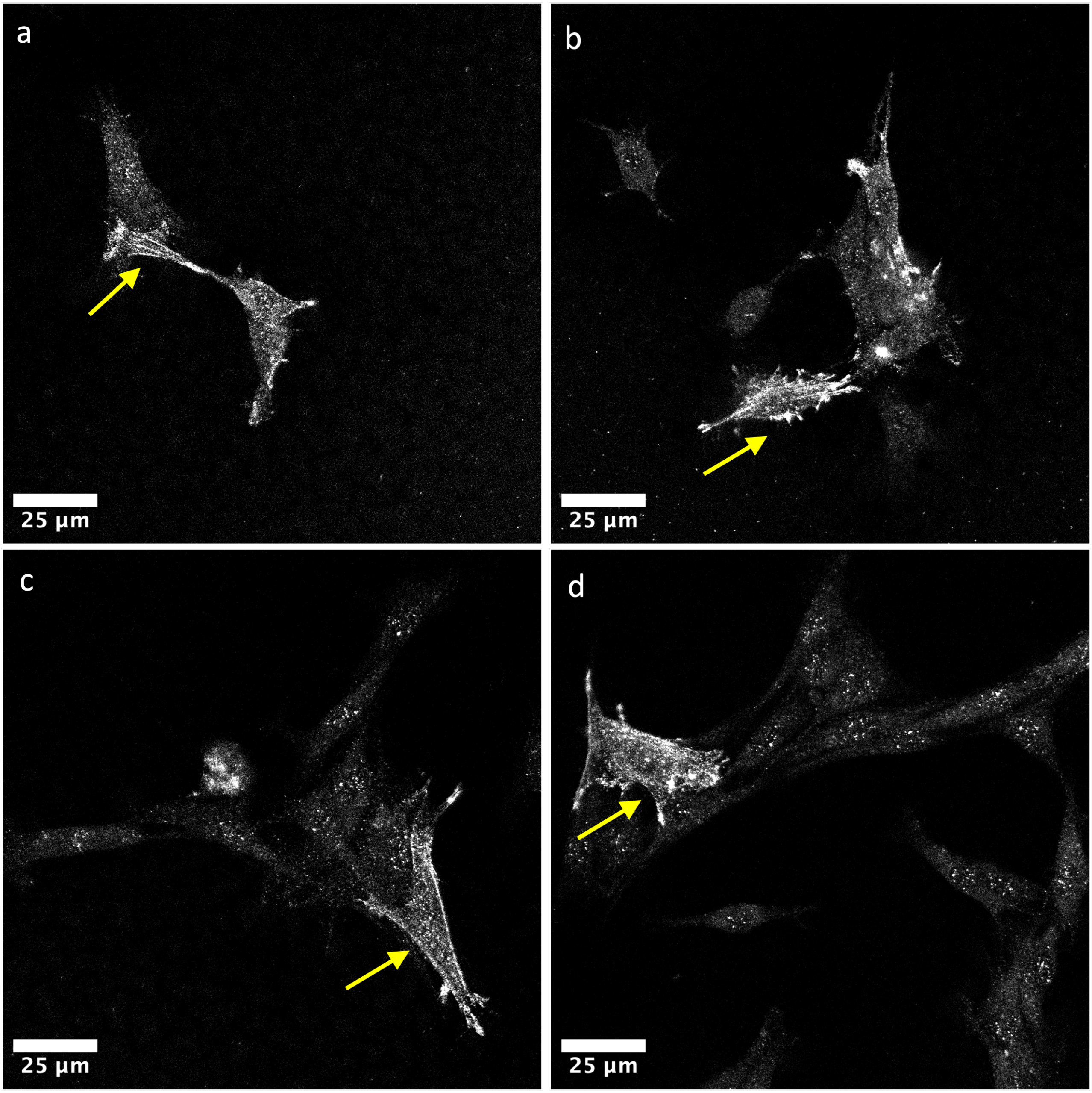


Fig. S7: **Spatial distribution of $\alpha$-SMA expression in mesangial cell clusters on fibrotic GelMA matrices.** Confocal immunofluorescence images (a–d) show expression of $\alpha$-smooth muscle actin ($\alpha$-SMA) in mesangial cells cultured on 10% GelMA hydrogels (fibrotic matrix) under chemical control conditions (absence of high glucose and TGF-$\beta$). Yellow arrows highlight the "leader" cell within each cluster, which consistently exhibits the highest $\alpha$-SMA expression regardless of the size of the associated cluster. (a–b) In smaller clusters, non-leader cells display moderate levels of $\alpha$-SMA, indicating partial contractile activity across the cluster. (c–d) In larger clusters, $\alpha$-SMA expression is almost exclusively localized to the leader cell, while follower cells show minimal signal. This differential distribution suggests a mechanical load-balancing mechanism in fibrotic environments, where contractile responsibility is centralized in a single cell to protect the rest of the cluster from stress-induced strain or energy expenditure.

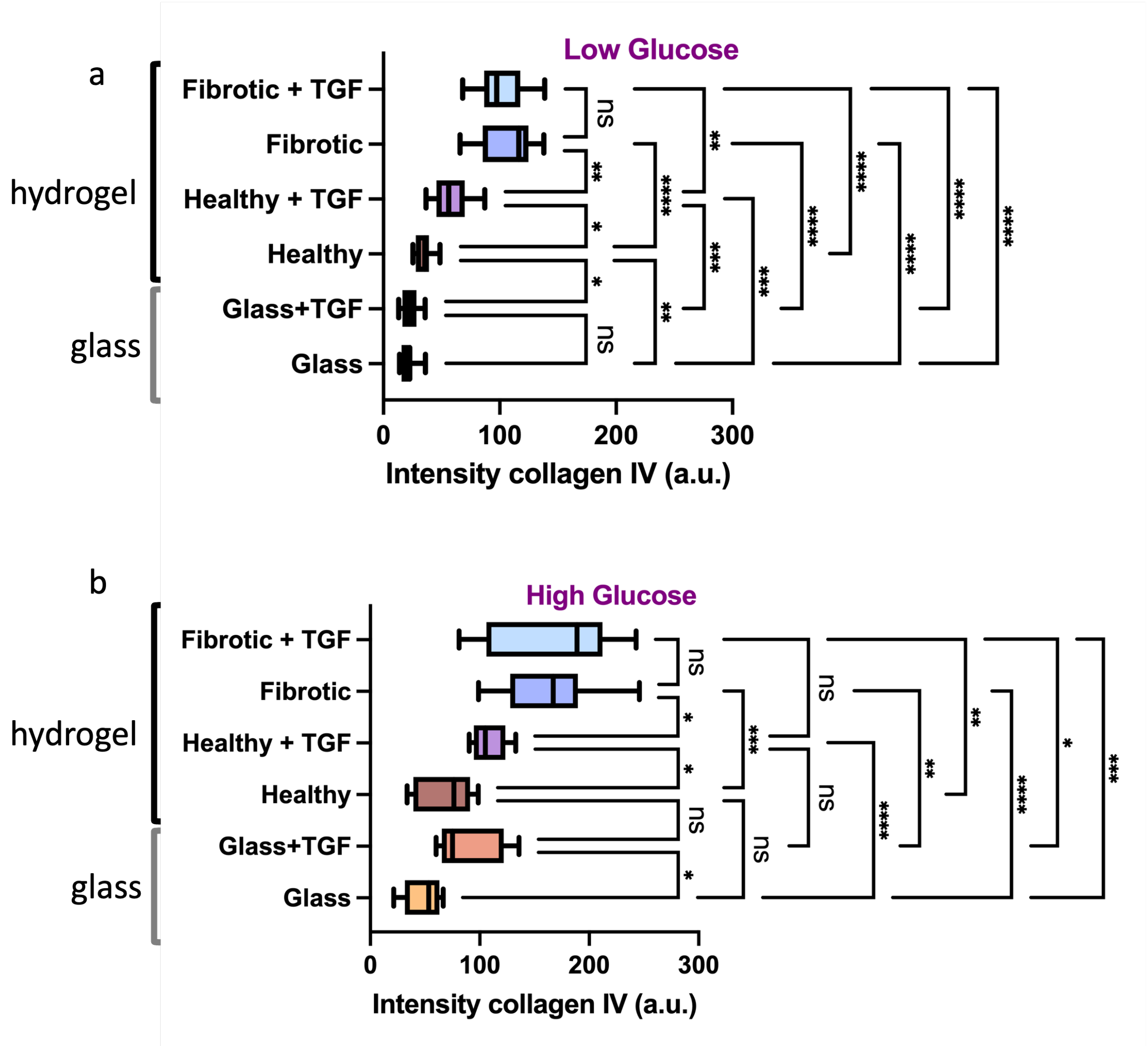


Fig. S8: **Collagen IV expression in mesangial cells across substrate stiffness and chemical conditions**. Quantification of collagen IV intensity (a.u.) in mesangial cells cultured under low glucose (top panel) and high glucose (bottom panel) conditions across different substrates and matrix configurations. (A) In the low glucose condition, the stiff glass substrate (50–80 GPa) fails to induce significant collagen IV expression, whereas softer 3D hydrogel matrices—healthy (5% GelMA, ∼2.5 kPa) and fibrotic (10% GelMA, ∼30 kPa)—show progressively higher expression, especially when coupled with TGF-$\beta$. This demonstrates the importance of matrix stiffness and 3D architecture, beyond simple substrate rigidity, in regulating ECM biosynthesis.(B) Under high glucose conditions, collagen IV expression is upregulated across all substrates. However, even with TGF-$\beta$, 2D glass fails to match the elevated levels observed in the fibrotic matrix, highlighting that traditional 2D cultures may significantly under-represent fibrotic phenotypes. These results support the necessity of 3D, stiffness-tuned environments with bioactive cues (e.g., RGD motifs) to accurately study fibrogenic responses of mesangial cells. Statistical significance was calculated using one-way ANOVA with multiple comparisons: $*p < 0.05$; $**p < 0.01$; $***p < 0.001$; $****p < 0.0001$; ns = not significant.

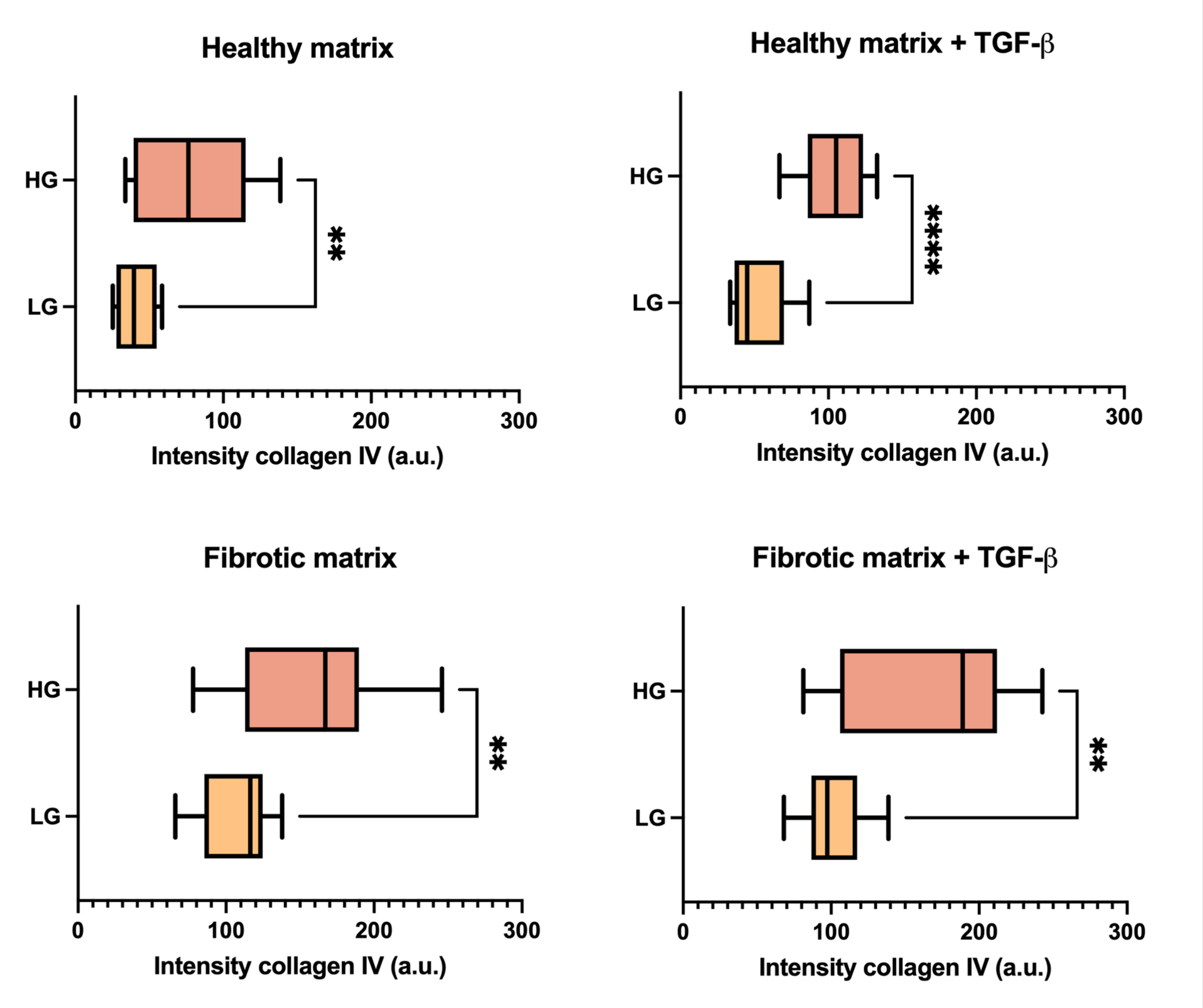


Fig. S9: **Effect of glucose concentration on collagen IV expression across mechanochemical conditions.**. Collagen IV intensity in mesangial cells is compared between high glucose (HG) and low glucose (LG) conditions across four matrix environments: healthy GelMA matrix (top left), healthy matrix + TGF-$\beta$ (top right), fibrotic GelMA matrix (bottom left), and fibrotic matrix + TGF-$\beta$ (bottom right). In all cases, high glucose significantly increases collagen IV expression, a key component of the glomerular basement membrane and mesangial matrix. These results underscore the potent influence of glucose concentration on ECM production, independent of stiffness or inflammatory stimulation. Statistical significance: **$p < 0.01$; ****$p < 0.0001$.

#### Log-likelihood and Probabilistic Modeling

A dataset of severety measurements $\mathbf{Y}(\mathbf{X})$, $\mathbf{Y} \in \mathbb{R}^N$, can be modelled as a probability density function $P(\mathbf{Y}|\mathbf{X}, \theta)$ of variables $\theta \in \mathbb{R}^n$ and samples $\mathbf{Y} = [y_1, y_2, ..., y_N]$ $(y_i \geq 0)$. Here, $\mathbf{X} = [\mathbf{x}_1, \mathbf{x}_2, ..., \mathbf{x}_N]$, $\mathbf{x}_i \in \mathbb{R}^3$ are 3D entry points representing glucose level, TGF, and matrix condition. As a result, every dataset can be expressed as a likelihood function

$$\mathcal{L}(\mathbf{X}, \theta|\mathbf{Y}) = \prod_{i=1}^{n} P(y_i|\mathbf{x}_i, \theta) \tag{1}$$

that calculates the probability of observing a vector of severety $\mathbf{Y}(\mathbf{X})$ from random draws of the distribution $P$ defined by the variables $\xi$ as a joint probability distribution. Thus, the dataset can be modeled with the Eq. 1 that needs to be estimated.

Assuming that $P(y_i|\mathbf{x}_i, \theta)$ are Gaussian distributions of variables $\theta = [\mu, \sigma^2]$

$$P(y_i; \mathbf{x}_i, \mu(\mathbf{x}_i), \sigma^2) = \frac{1}{\sqrt{2\pi\sigma^2}} exp\Big( - \frac{1}{2} \frac{(y_i - \mu(\mathbf{x}_i))^2}{\sigma^2} \Big) \tag{2}$$

the likelihood function converts to a new Gaussian distribution

$$\mathcal{L}(\mathbf{X}, \mu, \sigma^2|\mathbf{Y}) = \frac{1}{\sqrt{2\pi\sigma^2}} exp\Big( - \frac{1}{2\sigma^2} \sum_{i=1}^{N} (y_i - \mu(\mathbf{x}_i))^2 \Big) \tag{3}$$

Thus the modeling problem is reduced to the optimization of $\mathcal{L}(\mathbf{X}|\xi)$ to find the means and the standard deviation $\mu$ and $\sigma^2$.

In this optimization problem, we only need to find the optimal variables for which the likelihood is maximum. Thus, instead of directly maximizing the Eq.3, it is convenient to maximize its logarithm to remove the exponentiation.

$$l(\mathbf{X}, \mu, \sigma^2|\mathbf{Y}) = \ln(\mathcal{L}(\mathbf{X}, \mu, \sigma^2|\mathbf{Y})) = -\frac{N}{2} ln(2\pi) - \frac{N}{2} ln(\sigma^2) - \frac{1}{2\sigma^2} \sum_{i=1}^{N} (y_i - \mu(\mathbf{x}_i))^2 \tag{4}$$

This is the so-called log-likelihood equation. This modeling aims to find a predictive function based on the measurements for any entry value $\mathbf{x}$. In other words, we are seeking a $y^*(\mathbf{x}; \theta^*)$ that is built based on the measured data. $y^*$ can be expressed as a simple line function or any sort of mathematical function. In any case, Eq. 4 allows the optimization of a hypothetical function $y^*$ by adding it to $\mu$ as a constraint. This means we can optimize

$$l(\mathbf{X}, \mu, \sigma^2, \theta^*|\mathbf{Y}) = \ln(\mathcal{L}(\mathbf{X}, \mu, \sigma^2, \theta^*|\mathbf{Y})) = -\frac{N}{2} ln(2\pi) - \frac{N}{2} ln(\sigma^2) - \frac{1}{2\sigma^2} \sum_{i=1}^{N} (y_i - y^*(\mathbf{x}_i; \theta^*))^2 \tag{5}$$

where $y^*$ has a bias that encapsulates $\mu$ as well. having this definition, the optimization problem becomes a linear regression problem by means of log-likelihood as the sampling probability model. The targets would be finding the optimal parameters $\theta^*$ and a standard deviation $\sigma^2$ to represent an average error.

#### Linear Regression and Gaussian Process

The linear regression problem presented in the previous section can be defined based on any possible function that may explain the behavior of data. In most real-world problems, the true function is unknown or nonexistent, and we must make assumptions. Severity in relation to organic quantities, such as cellular matrix conditions, is an organic quantity with a nonlinear relation. There is no basis for making assumptions about its behavior, and it is impossible to model precisely by a linear function. Furthermore, in this study, the datasets are extremely undersampled and lack intermediate measurements.

The data analysis problem here reduces to finding functions that fit with separated collections of points. Examples can be seen in Fig.??. We treat this problem in two ways: 1. as a linear 1D curve fitting problem where only the trend line is important, and 2. as a generic linear 1D or N-dimensional problem where the shape of the function $y^*$ is unknown. In both cases, we assume that a linear function exists that explains the severity of the behavior with a certain degree of accuracy.

A line function can be optimized to investigate the trend line between two collections of points. In this case

$$y^*(\mathbf{x}_i; m, b) = m\mathbf{x}_i + b \tag{6}$$

where $\theta^* = [m, b]$ is the set of optimizable variables to define a line that is supposed to fit the severity trend.

This assessment of Eq.5 can be conducted purely statistically without any hypothesis about the model $y^*$. Gaussian Process (GP) is a stochastic process that is used to find a joint probability distribution of the data by fitting Gaussian distributions to every collection of samples from the dataset [**Rasmussen˙Williams˙2006**]. In other words, GP finds the distribution of all functions that fit with the dataset. Thus, the dataset is assumed to fit with an unknown function $f$ with an error $\epsilon$

$$y_i = f(\mathbf{x}_i) + \epsilon = (\mathbf{X}^T\mathbf{w})_i + \epsilon \tag{7}$$

where the error is assumed to be

$$\epsilon \sim \mathcal{N}(0, \sigma^2) \tag{8}$$

Assuming that the priors $\mathbf{w}$ satisfy a Gaussian distribution, and using Bayes' rule, it can be shown that a Gaussian distribution can be inferred from which the noisy observations $\mathbf{Y}$ are pulled. The standard deviation of this Gaussian distribution is specified by covariances of observation variables $\mathbf{X}$ such that for a zero-meaned measurement set $\mathbf{Y}$, we can write [**Rasmussen˙Williams˙2006**]

$$\mathbf{Y} \sim \mathcal{N}(0, K(\mathbf{X}, \mathbf{X}) + \sigma^2\mathbf{I}) \tag{9}$$

where $K$ is known as the kernel function and calculates the covariance of any pair of variables.

This formalism allows us to explain the observations using a probability density function of all possible functions fitting with the dataset instead of finding an explicit model $f$. We can use Eq.9 to identify a predictive model for a set of test points $\mathbf{X}*$, and we would have [**Rasmussen˙Williams˙2006**]

$$\mathbf{Y}* \sim \mathcal{N}(0, K(\mathbf{X}*, \mathbf{X}*)) \tag{10}$$

and one can show the relation

$$\begin{pmatrix} \mathbf{Y} \\ \mathbf{Y}* \end{pmatrix} \sim \mathcal{N}\left(0, \begin{bmatrix} K(\mathbf{X}, \mathbf{X}) + \sigma^2\mathbf{I} & K(\mathbf{X}, \mathbf{X}*) \\ K(\mathbf{X}*, \mathbf{X}) & K(\mathbf{X}*, \mathbf{X}*) \end{bmatrix}\right) \tag{11}$$

This is the so-called Gaussian Process, which begins by modeling the measurement noise with a Gaussian function and derives a posterior predictive probability distribution of all possible observations given the available dataset, as shown in Eq.11. This model-less regression method allows us to identify a generic curve that fits with the data.

The covariance function is the only user-defined component of the GP method. We chose to use one of the most common kernels, known as the Radial Basis Function (RBF), for data processing tasks that involved GP. RBF is a squared exponential function of the distance between variables and for every pair of point vectors $\mathbf{X}_1$ and $\mathbf{X}_2$, it is given by

$$K(\mathbf{X}_1, \mathbf{X}_2) = exp\left(-\frac{d(\mathbf{X}_1, \mathbf{X}_2)}{2l^2}\right) \tag{12}$$

where $d$ calculates the Euclidian distance between each pair of points within the point vectors and $l$ is a scaling function, generally known as length scale. The length scale determines the smoothness of the functions specified by the inferred distribution. The linear regression problem using maximum likelihood and GP schemes and optimizing the kernel hyperparameters are covered in the next section.

#### Optimization Methods

For each cell condition, the variance of severity is significant. Besides, the dataset is severely undersampled with respect to the cell conditions. This situation simply causes the GP to overfit a distribution with the measurements. To avoid this issue, in the MCMC process, a slight Gaussian noise is added to the data points to vary their severity values and their corresponding cell variables on each step. Data augmentation is used to artificially expand the dataset and to force the optimization to converge to a more general model. This is achieved by applying a slight noise to the data points at each sample draw. The additive noise is Gaussian with the variance below

$$\sigma_x^2 = \frac{0.1}{e}, \quad \sigma_y^2(x) = 0.1\sigma_d^2(x) \tag{13}$$

where $\sigma_x^2$, and $\sigma_y^2$ and the variance of the Gaussian noise in the x and y directions, respectively, and $e$ is the Euler's number. In this definition, $\sigma_d^2(x)$ is the variance of the dataset for a corresponding $x$ value, making $\sigma_y^2$ a function of $x$. Since the dataset is severely undersampled in the $x$ axis, $\sigma_x^2$ is chosen to be a relatively large value.

The optimizable variables are also specified by the model and the log-likelihood equation expressed in Eq.5. For instance, for a line fitting problem, the line function parameters plus the logarithm of the variance of the Gaussian noise function (Eq.2) are variables of interest. We specify the last variable as $log(f)$ where $f := \sigma^2$. Similarly, to optimize a curve using the Gaussian process, the line variables will be substituted with the covariance function parameters. The RBF function defined at Eq.12 has one variable, which is the length scale parameter. This leaves a set of two parameters to be optimized using the MCMC approach, $\{l, log(f)\}$, for linear regression using the GP method. Thus, the optimization can be conducted using the MCMC approach.

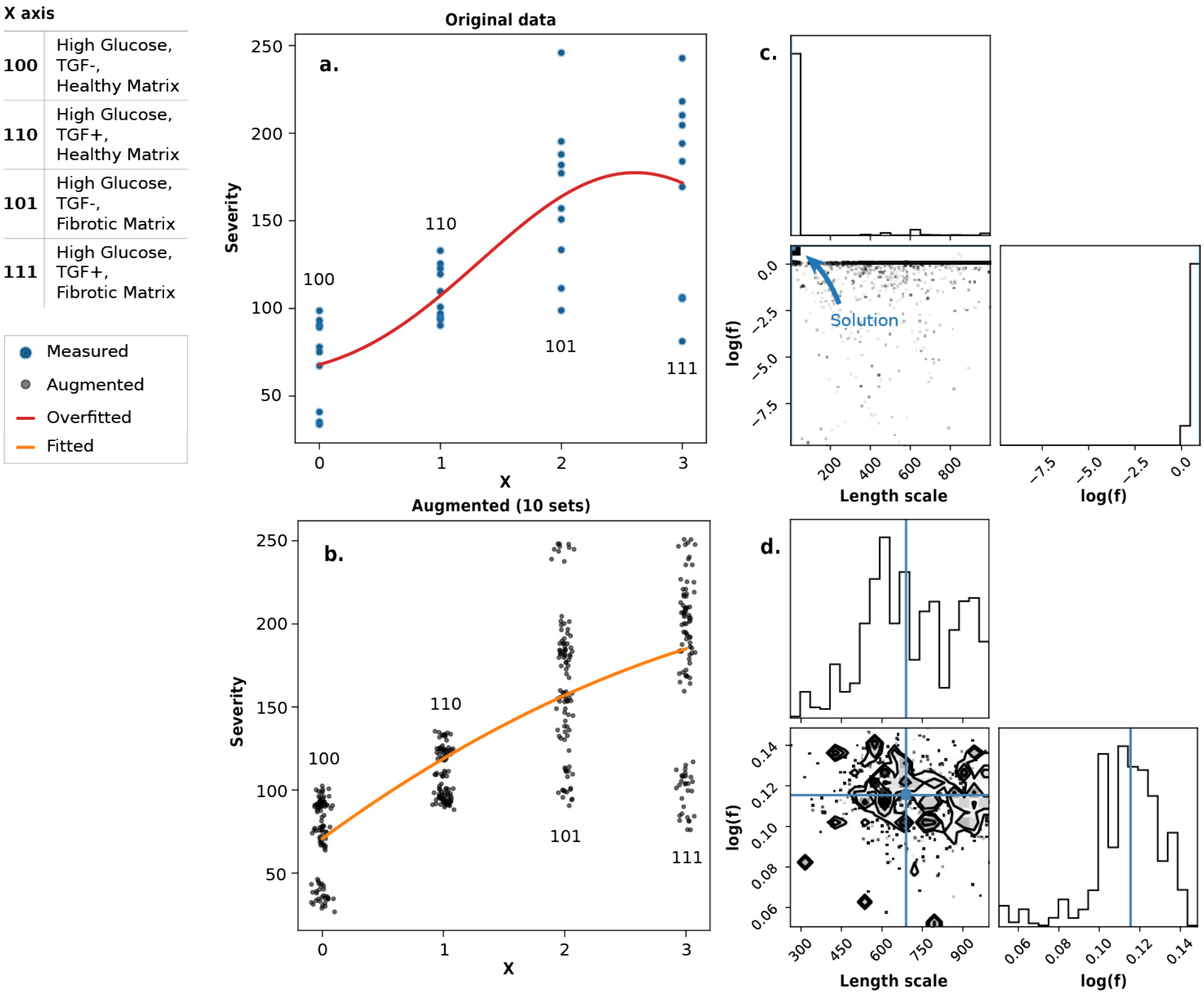


Fig. S10: **Data augmentation to address the undersampling errors**. **a.** the severity measurements of ColIV in high glucose conditions and the fitted curve. Since the dataset is undersampled in the $x$-axis, the optimized curve is overfitted. **b.** the severity measurements of ColIV in high glucose conditions and the fitted curve when the data that is augmented with Gaussian noise. **c.** The estimated posterior probability density function that is evaluated by the MCMC approach in 20000 steps. It is evident that the solution is highly localized at the minimum of the search range at every dimension, which ended up in the overfitted curve in **a**. **d.** The posterior probability density function is given the data points at each step. The estimated distribution has a new maximum which is away from minimums.

Figure S10 depicts the optimization results for both unaugmented and augmented data points using the GP method. In Fig.S10**a** and Fig.S10**b**, 10 sets of samples are depicted that are given to the MCMC optimizer. The curves are plotted using the optimized parameters in each case. The corner plots illustrated in Fig.S1**c** and Fig.S10**d** show the estimated posterior probability density functions from which the solutions can be found. As can be deduced from Fig.S10**a** and Fig.S10**c**, the problem, by default, is too underdetermined in the x-axis and badly conditioned, which results in an overfitted curve. This issue is well addressed by adding random noise to each sample that is drawn from the dataset. Figures S10**a** and S10**c** show that the posterior probability density function and the equivalent optimal curve are broadened and generalized and are a more suitable predictive model.

Similarly, the same analysis can be performed on the 3D predictive model problem. Figure S11 shares insight into the log-likelihood of the length scale parameter for 20000 samples and its distribution. The distribution

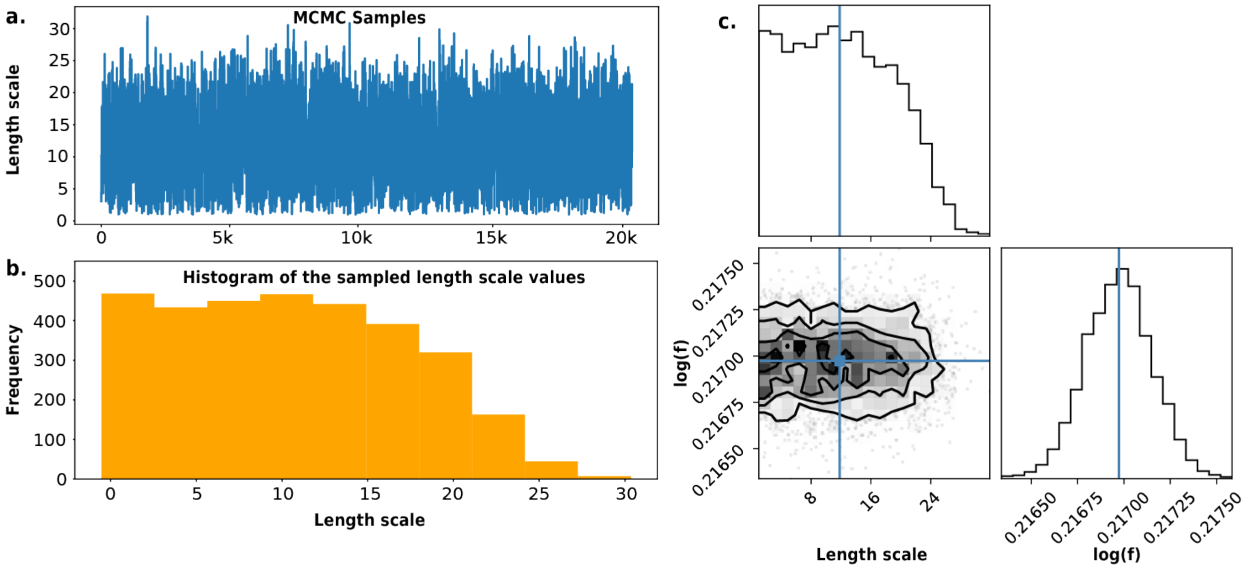


Fig. S11: **Posterior probability density function of the 3D predictive model obtained by the GP method.a.** 20000 length scale samples drawn in the MCMC process during the random walk and evolution steps. These values are given to the RBF to calculate the corresponding log-likelihoods. **b.** The histogram of **a** depicting the probability distribution of the results with the highest likelihood. **c.** The probability density function of both the optimizable parameters and the discovered solution.

can be observed in the histogram plot shown in Fig.S11**b**. The estimated posterior probability density function is shown as a corner plot in Fig.S11**c**.

In all cases, the final solution was determined by averaging the samples corresponding to the 50 percent highest log-likelihood.

### Supplementary Tables

**Statistical analysis:** Two-way ANOVA with multiple comparisons. Sample size: n=38 per group. Significance level $\alpha$=0.05.

Table 1: **Supplementary Table 1**: 2-way ANOVA of $\alpha$-SMA expressions (suppl. to Fig3e)

| Comparison | Mean Diff (95% CI) | Summary | Adjusted P Value |
|---|---|---|---|
| LG:Control vs. LG:Healthy matrix | -722.2 (-1404 to -40.12) | * | **0.0272** |
| LG:Control vs. LG:Fibrotic matrix | -1770 (-2327 to -1214) | **** | **<0.0001** |
| LG:Control vs. LG+:Control | -153.9 (-836.1 to 528.2) | ns | 0.9999 |
| LG:Control vs. LG+:Healthy matrix | -1642 (-2324 to -959.4) | **** | **<0.0001** |
| LG:Control vs. LG+:Fibrotic matrix | -1926 (-2501 to -1350) | **** | **<0.0001** |
| LG:Control vs. HG:Control | 17.65 (-664.5 to 699.8) | ns | >0.9999 |
| LG:Control vs. HG:Healthy matrix | -1074 (-1621 to -527.0) | **** | **<0.0001** |
| LG:Control vs. HG:Fibrotic matrix | -2348 (-2895 to -1801) | **** | **<0.0001** |
| LG:Control vs. HG+:Control | -571.9 (-1254 to 110.3) | ns | 0.2036 |
| LG:Control vs. HG+:Healthy matrix | -1737 (-2323 to -1151) | **** | **<0.0001** |
| LG:Control vs. HG+:Fibrotic matrix | -2797 (-3399 to -2195) | **** | **<0.0001** |
| LG:Healthy matrix vs. LG:Fibrotic matrix | -1048 (-1605 to -491.3) | **** | **<0.0001** |
| LG:Healthy matrix vs. LG+:Control | 568.3 (-113.8 to 1250) | ns | 0.2115 |
| LG:Healthy matrix vs. LG+:Healthy matrix | -919.3 (-1601 to -237.2) | *** | **0.0007** |
| LG:Healthy matrix vs. LG+:Fibrotic matrix | -1203 (-1779 to -627.5) | **** | **<0.0001** |
| LG:Healthy matrix vs. HG:Control | 739.9 (57.77 to 1422) | * | **0.0205** |
| LG:Healthy matrix vs. HG:Healthy matrix | -351.7 (-898.6 to 195.2) | ns | 0.6137 |
| LG:Healthy matrix vs. HG:Fibrotic matrix | -1626 (-2172 to -1079) | **** | **<0.0001** |
| LG:Healthy matrix vs. HG+:Control | 150.4 (-531.7 to 832.5) | ns | 0.9999 |
| LG:Healthy matrix vs. HG+:Healthy matrix | -1015 (-1601 to -429.0) | **** | **<0.0001** |
| LG:Healthy matrix vs. HG+:Fibrotic matrix | -2075 (-2676 to -1473) | **** | **<0.0001** |
| LG:Fibrotic matrix vs. LG+:Control | 1617 (1060 to 2173) | **** | **<0.0001** |
| LG:Fibrotic matrix vs. LG+:Healthy matrix | 128.9 (-428.0 to 685.9) | ns | 0.9998 |
| LG:Fibrotic matrix vs. LG+:Fibrotic matrix | -155.1 (-575.3 to 265.1) | ns | 0.9878 |
| LG:Fibrotic matrix vs. HG:Control | 1788 (1231 to 2345) | **** | **<0.0001** |
| LG:Fibrotic matrix vs. HG:Healthy matrix | 696.6 (317.1 to 1076) | **** | **<0.0001** |
| LG:Fibrotic matrix vs. HG:Fibrotic matrix | -577.3 (-956.8 to -197.8) | *** | **0.0002** |
| LG:Fibrotic matrix vs. HG+:Control | 1199 (641.7 to 1756) | **** | **<0.0001** |
| LG:Fibrotic matrix vs. HG+:Healthy matrix | 33.19 (-400.8 to 467.2) | ns | >0.9999 |
| LG:Fibrotic matrix vs. HG+:Fibrotic matrix | -1027 (-1481 to -571.8) | **** | **<0.0001** |

**Statistical analysis:** Brown-Forsythe and Welch ANOVA tests. Sample size: n=35 per group. Significance level $\alpha$=0.05.

Table 2: **Supplementary Table 2**: ANOVA tests of $\alpha$-SMA booleanized (suppl. to Fig3k)

| Comparison | Mean Diff (95% CI) | Summary | Adjusted P Value |
|---|---|---|---|
| 1 vs. 2 | -9.785 (-20.00 to 0.4277) | ns | 0.0717 |
| 1 vs. 3 | -38.63 (-53.61 to -23.65) | **** | **<0.0001** |
| 1 vs. 4 | -42.65 (-58.04 to -27.26) | **** | **<0.0001** |
| 1 vs. 5 | -44.05 (-59.90 to -28.19) | **** | **<0.0001** |
| 1 vs. 6 | -47.64 (-60.49 to -34.80) | **** | **<0.0001** |
| 1 vs. 7 | -68.30 (-83.85 to -52.76) | **** | **<0.0001** |
| 1 vs. 8 | -87.18 (-113.2 to -61.16) | **** | **<0.0001** |
| 2 vs. 3 | -28.84 (-42.60 to -15.09) | **** | **<0.0001** |
| 2 vs. 4 | -32.87 (-46.99 to -18.75) | **** | **<0.0001** |
| 2 vs. 5 | -34.26 (-48.88 to -19.64) | **** | **<0.0001** |
| 2 vs. 6 | -37.86 (-49.01 to -26.71) | **** | **<0.0001** |
| 2 vs. 7 | -58.52 (-72.78 to -44.26) | **** | **<0.0001** |
| 2 vs. 8 | -77.39 (-102.8 to -52.00) | **** | **<0.0001** |
| 3 vs. 4 | -4.024 (-21.69 to 13.64) | ns | >0.9999 |
| 3 vs. 5 | -5.418 (-23.50 to 12.66) | ns | >0.9999 |
| 3 vs. 6 | -9.016 (-24.66 to 6.624) | ns | 0.7921 |
| 3 vs. 7 | -29.68 (-47.50 to -11.86) | **** | **<0.0001** |
| 3 vs. 8 | -48.55 (-75.79 to -21.32) | **** | **<0.0001** |
| 4 vs. 5 | -1.395 (NA) | ns | >0.9999 |
| 4 vs. 6 | -4.992 (-21.09 to 11.10) | ns | >0.9999 |
| 4 vs. 7 | -25.65 (-43.95 to -7.354) | *** | **0.0005** |
| 4 vs. 8 | -44.53 (-72.12 to -16.94) | **** | **<0.0001** |
| 5 vs. 6 | -3.598 (-20.16 to 12.96) | ns | >0.9999 |
| 5 vs. 7 | -24.26 (-42.99 to -5.531) | ** | **0.0019** |
| 5 vs. 8 | -43.13 (-70.99 to -15.28) | *** | **0.0001** |
| 6 vs. 7 | -20.66 (-36.92 to -4.400) | ** | **0.0026** |
| 6 vs. 8 | -39.53 (-65.99 to -13.08) | *** | **0.0003** |
| 7 vs. 8 | -18.87 (-46.60 to 8.850) | ns | 0.5419 |

**Note:** 1 = 000 (no diabetes, no TGF, normal stiffness), 2 = 100 (diabetes only), 3 = 010 (TGF only), 4 = 110 (diabetes + TGF), 5 = 001 (stiffness only), 6 = 011 (TGF + stiffness), 7 = 101 (diabetes + stiffness), 8 = 111 (diabetes + TGF + stiffness).

**Statistical analysis:** Brown-Forsythe and Welch ANOVA tests. Sample size: n=20 per group. Significance level $\alpha$=0.05.

Table 3: **Supplementary Table 3**: ANOVA tests of Col-IV booleanized (suppl. to Fig3l)

| Comparison | Mean Diff (95% CI) | Summary | Adjusted P Value |
|---|---|---|---|
| 1 vs. 2 | -35.05 (-67.85 to -2.255) | * | **0.0333** |
| 1 vs. 3 | -22.59 (-42.93 to -2.252) | * | **0.0239** |
| 1 vs. 4 | -73.60 (-93.93 to -53.28) | **** | **<0.0001** |
| 1 vs. 5 | -71.35 (-104.9 to -37.82) | *** | **0.0002** |
| 1 vs. 6 | -66.67 (-96.12 to -37.22) | **** | **<0.0001** |
| 1 vs. 7 | -128.8 (-185.8 to -71.87) | *** | **0.0001** |
| 1 vs. 8 | -136.5 (-208.5 to -64.54) | *** | **0.0006** |
| 2 vs. 3 | 12.46 (-21.73 to 46.65) | ns | 0.9824 |
| 2 vs. 4 | -38.55 (-72.73 to -4.374) | * | **0.0202** |
| 2 vs. 5 | -36.29 (-76.79 to 4.202) | ns | 0.1063 |
| 2 vs. 6 | -31.62 (-69.88 to 6.649) | ns | 0.1721 |
| 2 vs. 7 | -93.79 (-152.7 to -34.85) | *** | **0.0009** |
| 2 vs. 8 | -101.4 (-173.6 to -29.29) | ** | **0.0035** |
| 3 vs. 4 | -51.01 (-75.76 to -26.27) | **** | **<0.0001** |
| 3 vs. 5 | -48.75 (-83.54 to -13.97) | ** | **0.0029** |
| 3 vs. 6 | -44.08 (-75.76 to -12.39) | ** | **0.0029** |
| 3 vs. 7 | -106.3 (-163.1 to -49.41) | *** | **0.0004** |
| 3 vs. 8 | -113.9 (-186.1 to -41.74) | ** | **0.0020** |
| 4 vs. 5 | 2.258 (NA) | ns | >0.9999 |
| 4 vs. 6 | 6.936 (-24.74 to 38.61) | ns | >0.9999 |
| 4 vs. 7 | -55.24 (-112.1 to 1.595) | ns | 0.0595 |
| 4 vs. 8 | -62.90 (-135.1 to 9.266) | ns | 0.1073 |
| 5 vs. 6 | 4.677 (-34.10 to 43.45) | ns | >0.9999 |
| 5 vs. 7 | -57.50 (-116.1 to 1.113) | ns | 0.0571 |
| 5 vs. 8 | -65.15 (-137.6 to 7.308) | ns | 0.0971 |
| 6 vs. 7 | -62.18 (-119.8 to -4.502) | * | **0.0292** |
| 6 vs. 8 | -69.83 (-142.0 to 2.387) | ns | 0.0618 |
| 7 vs. 8 | -7.656 (-87.69 to 72.38) | ns | >0.9999 |

**Note:** 1 = 000 (no diabetes, no TGF, normal stiffness), 2 = 100 (diabetes only), 3 = 010 (TGF only), 4 = 110 (diabetes + TGF), 5 = 001 (stiffness only), 6 = 011 (TGF + stiffness), 7 = 101 (diabetes + stiffness), 8 = 111 (diabetes + TGF + stiffness).
